# *NCOA3* identified as a new candidate to explain autosomal dominant progressive hearing loss

**DOI:** 10.1101/2020.06.07.138909

**Authors:** Rodrigo Salazar da Silva, Vitor Lima Goes Dantas, Leandro Ucela Alves, Ana Carla Batissoco, Jeanne Oiticica, Elizabeth A Lawrence, Abdelwahab Kawafi, Yushi Yang, Fernanda Stávale Nicastro, Beatriz Caiuby Novaes, Chrissy Hammond, Erika Kague, Regina Célia Mingroni Netto

## Abstract

Hearing loss is a frequent sensory impairment in humans and genetic factors account for an elevated fraction of the cases. We have investigated a large family of five generations, with 15 reported individuals presenting non-syndromic, sensorineural, bilateral and progressive hearing loss, segregating as an autosomal dominant condition. Linkage analysis, using SNP-array and selected microsatellites, identified a region of 13cM in chromosome 20 as the best candidate to harbour the causative mutation. After exome sequencing and filtering of variants, only one predicted deleterious variant in the *NCOA3* gene (NM_181659, c.2810C>G; p.Ser937Cys) fit in with our linkage data. RT-PCR, immunostaining and *in situ* hybridization showed expression of *ncoa3* in the inner ear of mice and zebrafish. We generated a stable homozygous zebrafish mutant line using the CRISPR/Cas9 system. *ncoa3−/−* did not display any major morphological abnormalities in the ear, however, anterior macular hair cells showed altered orientation. Surprisingly, chondrocytes forming the ear cartilage showed abnormal behaviour in *ncoa3−/−*, detaching from their location, invading the ear canal and blocking the cristae. Adult mutants displayed accumulation of denser material wrapping the otoliths of *ncoa3−/−* and increased bone mineral density. Altered zebrafish swimming behaviour corroborates a potential role of *ncoa3* in hearing loss. In conclusion, we identified a potential candidate gene to explain hereditary hearing loss, and our functional analyses suggest subtle and abnormal skeletal behaviour as mechanisms involved in the pathogenesis of progressive sensory function impairment.

## Introduction

Hearing loss affects almost 466 million people worldwide and is estimated to affect more than 900 million people by 2050 (1). Genetic factors play an important role in the pathogenesis of the disease, with up to 55% of age-related hearing loss attributed to genetics (2). Approximately 70% of hereditary deafness cases are non-syndromic (3), of which 20% are autosomal dominant (2). Autosomal dominant non-syndromic hearing loss (ADNSHL) is typically progressive with late and variable average age of onset, which depends on the nature of the type of mutation and affected gene.

Mapping studies of large families have contributed to the identification of several genes associated with hearing loss (4). Recently, whole-genome and exome sequencing, in combination with familial cases, have boosted the identification of causal genes (4–8). The genetic complexity of the condition is highlighted by the large number of genes identified as associated to monogenic inheritance of non-syndromic hearing loss (~130), and those related to ADNSHL (~50) (9). New genes are still to be identified, however, given the extensive genetic heterogeneity underpinning the origin of hearing loss, newly identified genes and variants are rarely found, in only one or a few pedigrees, making their confirmation by reproducibility a challenging task. Therefore, functional studies are key to validate the genetic findings.

Hearing loss associated genes fall into common categories such as maintenance of ionic homeostasis, formation of hair cell stereocilia and regulation of gene transcription (10–15). Recently, other pathways have also been suggested to play a role in disease pathogenesis; such as collagen biogenesis and homeostasis (16–18). Thus, the identification of novel candidate genes associated with hearing loss could reveal new molecular players involved in the condition and potential therapeutics.

Here, we describe a large Brazilian family in which hearing loss segregates as an autosomal dominant trait. By linkage analysis and exome sequencing we identified a rare missense variant in the gene *NCOA3* (NM_181659:c.2810C>G:p.Ser937Cys) that segregated in the pedigree with hearing loss. We detected expression of the gene in mice and zebrafish ears. Using CRISPR/Cas9 genome editing, we generated a zebrafish *ncoa3−/−* which showed cartilage behaviour abnormalities in the larval sensorial region of the ear, amorphous material accumulation in proximity with adult otoliths, higher mineral density and abnormal adult swimming behaviour. Our work provides evidence of *NCOA3* playing an important role in skeletal system homeostasis and suggests *NCOA3* as a potential candidate gene associated with hearing impairment.

## Results

### Clinical findings in patients of a family with autosomal dominant, non-syndromic, sensorineural hearing loss

The five-generation Brazilian family examined in this study presented individuals affected by nonsyndromic, progressive, sensorineural, bilateral, moderate-to-profound hearing loss, segregating as an autosomal dominant condition (Figure 1A). Affected and non-affected individuals were submitted to audiological tests (Figure 1B). Age of onset of hearing loss varied from 4 to 35 years, with the average age of onset being 12 years old (Table 1).

**Figure 1.**
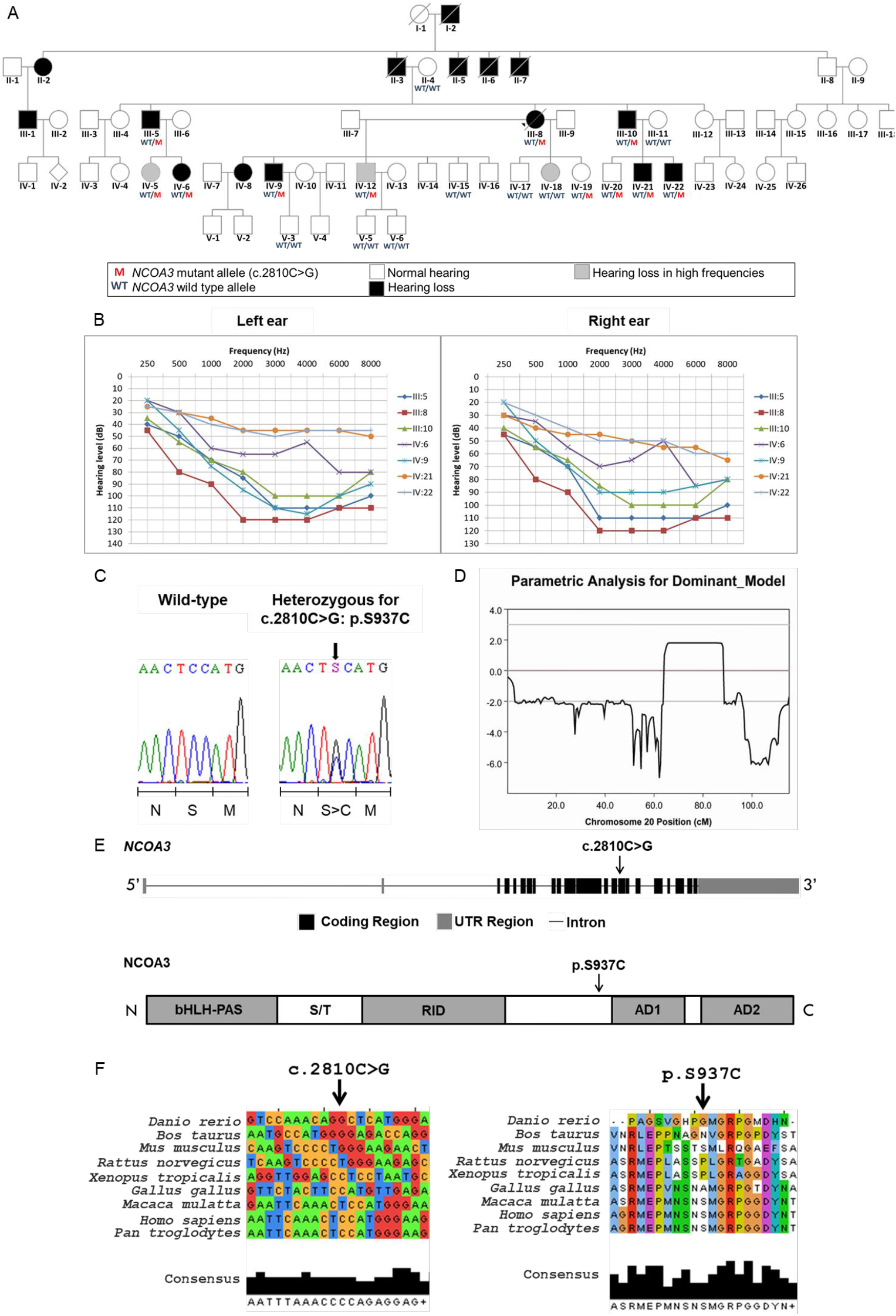
A rare variant in *NCOA3*, a gene which codes a nuclear receptor coactivator, segregates with hearing loss in the family. A) Pedigree showing the segregation of the *NCOA3* variant (NM_181659: c.2810C>G: p.Ser937Cys). B) 14 audiometric profiles (divided in right and left ear) of 7 patients affected with sensorineural and bilateral hearing loss. Hearing thresholds until 20dB are considered normal. C) Chromatograms showing partial sequence from affected patient compared to a wild-type sequence. Arrow indicates position of the *NCOA3* variant, while scale bar below indicates which amino acid is changed when the variant is present. D) Multipoint LOD scores calculated with Merlin software for chromosome 20, using data from SNP arrays, under assumption of complete penetrance K=1. E) Schematics of *NCOA3* gene and its respective protein. bHLH= basic helix-loop-helix domain; PAS= Per/ARNT/Sim homologous domain; S/T= serine/threonine-rich region; RID= receptor interaction domain containing multiple LXXLL motifs; AD1 and AD2= activation domains 1 and 2. F) Multiple alignment of *NCOA3* gene and its orthologous (left), as well as multiple alignment of the respective proteins (right). Arrow indicates position of *NCOA3* variant (NM_181659: c.2810C>G: p.Ser937Cys).

**Table 1.** Reported ages of onset for hearing loss and ages at the time of clinical examination.

| Patient ID | Classification | Severity of hearing loss | Age at examination (years) | Age of onset (years) |
| --- | --- | --- | --- | --- |
| III-5 | Affected | Moderate in right ear and severe in the left ear | 48 | 8 |
| III-8 | Affected | Profound | 40 | 35 |
| III-10 | Affected | Moderate | 45 | 20 |
| IV-6 | Affected | Mild | 24 | 7 |
| IV-9 | Affected | Mild to severe | 22 | 6 |
| IV-21 | Affected | Moderate in right ear and mild in left ear | 8 | 6 |
| IV-22 | Affected | Mild | 4 | 4 |
| II-4 | Not affected | - | - | - |
| IV-5 | Not affected | Threshold of 35dB only at 6K | 27 | - |
| IV-12 | Not affected | Threshold of 30dB only at 6K | 26 | - |
| IV-15 | Not affected | - | 20 | - |
| IV-17 | Not affected | - | 15 | - |
| IV-18 | Not affected | Threshold of 28dB only at 6K | 11 | - |
| IV-19 | Not affected | - | 8 | - |
| IV-20 | Not affected | - | 11 | - |
| V-3 | Not affected | - | 3 | - |
| V-5 | Not affected | - | 5 | - |
| V-6 | Not affected | - | 4 | - |

We performed ear, nose and throat (ENT) physical examinations. Patients III-5, III-8, III-10, IV-20 and IV-21, showed normal results, as well as normal computed tomography scan of temporal bones, magnetic resonance imaging of the inner ear and thyroid ultrasound. IV-20 and IV-21 had bilateral mild earlobe hypogenesis. IV-21 showed coloboma auris. Other minor clinical findings were also observed. IV-5, IV-6: bifid uvula at oropharynx cavity examination; III-8 and III-10: normal responses from the otoneurological evaluation, including electrooculography with caloric tests; and III-5: despite the absence of vestibular complaints, showed right idiopathic vestibular weakness on caloric test.

### Linkage analysis points to a region of 13.5Mb on chromosome 20

LOD score values from SNP-array analysis, assuming both complete (K= 1) and incomplete (K= 0.8 and K=0,64) penetrance suggested linkage to chromosome 20(chr20:40201234-53724200, 64 and 88 cM, GRCh37/hg19), with maximum positive value of 1.806 (K= 1; Figure 1D) and 1.805 (K= 0.8 and K= 0,64). This region has 13,5 Mb andcontains161 genes. We selected twelve microsatellite markers mapped along chromosome 20 (Supplementary Figure 1), which confirmed our SNP-array analysis pointing to a candidate region between 58 and 79 cM (complete penetrance, K= 1, maximum Lod score= 1.006) and between 56 and 83 cM (incomplete penetrance, K= 0.8 and K= 0,64, maximum Lod score=2.286). No other chromosomal region showed higher Lod scores than the ones obtained in chromosome 20. The maximum two-point LOD score value for this genealogy was simulated for complete (k=1) and incomplete (K= 0.8 and K= 0,64) penetrance resulting in 4.214, 3.580 and 3.145, respectively.

### Variant in *NCOA3* identified as candidate for hearing loss by whole-exome sequencing

We conducted whole-exome sequencing in samples from two of the affected individuals (III-8 and III-10) (Figure 1A); obtaining approximately 70M reads per sample (read average length of 99 bp, average coverage of 120X and 98% of target bases with more than 20 reads). We selected autosomal, exonic, heterozygous and nonsynonymous variants with Q >30 and coverage > 20, checked them against public variant databases and 66 control samples (sequenced simultaneously), and filtered for variants with frequencies lower than 0.01. A total of 162 variants shared by both samples were obtained (Table 2; Supplementary Table 1). From these variants, only the NM_181659: c.2810C>G: p.S937C in *NCOA3* gene matched the suggestive positive LOD score region mapped in the chromosome 20, as indicated by the linkage analysis.

**Table 2.** Steps of variant filtering after exome sequencing of samples from two affected individuals. *= 66 control samples that were sequenced in the same batch. **= 1000 genomes, NHLBI Exome Sequencing Project, Online Archive of Brazilian Mutations databases.

| **Filtration steps of exonic variants** | **# of remaining variants** |
| --- | --- |
| Heterozygous variants found in both patients analysed | 9197 |
| Exclusion of low-quality variants | 9193 |
| Exclusion of variants with f>1% in control-samples* | 553 |
| Exclusion of variants with f>1% in databases** | 350 |
| Considering only variants in autosomes | 349 |
| Exclusion of variants in hypervariable genes | 302 |
| Exclusion of synonymous variants | 162 |
| Considering only variants in chromosome 20 | 3 |
| Considering only variants in the positive Lod score region | 1 |

**Abbreviations**

Only two variants were detected in genes previously described as associated to hearing loss: NM_001258370: c.A1565G:p.Gln522Arg (*DIAPH3*) and NM_005709: c.G946C:p.Glu316Gln (*USH1C*). The variants in *DIAPH3* and *USH1C* were investigated in the pedigree by Sanger sequencing, and their segregation was not compatible with the segregation of hearing loss in the family (Supplementary Figure 2). Moreover, copy-number variation was excluded after array-CGH (Agilent Technologies, 180K).

*NCOA3* (Nuclear Receptor Coactivator 3) comprises 23 exons, encoding a protein of 1420 amino acids, with a suggested function in the regulation of gene transcription, mediated by nuclear receptors and it has never been reported to be associated with hearing loss. The variant c.2810C>G in exon 15 is predicted to result in a p.Ser937Cys amino acid substitution within a highly conserved region among primates (Figures 1E and F). This variant was predicted to be damaging using several prediction tools: SIFT showed a damaging score of 0.030, and Polyphen2, a score of 0.905. MutationTaster2 predicted that it is a disease-causing mutation (score of 0.845). The variant, rs142951578, within *NCOA3* has been reported with low frequency by GnomAD (0.0003465), NHLBI-ESP (0.000538), 1000 genomes (0.001) and was not described by ABraOM.

We investigated the segregation of NM_181659: c.2810C>G: p.Ser937Cys in *NCOA3* by Sanger sequencing in 19 samples. This variant was found to be present in heterozygosis in all seven affected individuals and in 4 non-affected ones (Figure 1A and 1C). These four heterozygous non-affected individuals are within the range of onset of hearing loss observed in the family (4-35 years, Table 1), therefore, it is possible that manifestation of hearing loss will occur later.

### *Ncoa3* is expressed in the developing mouse cochlea and zebrafish ear

*Ncoa3* expression in mice has been reported for ovary, testis, liver, skeletal muscle and adipose tissue (45–47), and transcriptome studies have suggested its expression in the ear (48,49), however this has been poorly characterised. To determine the temporal pattern of *Ncoa3* expression in the inner ear of mice we performed RT-PCR and immunofluorescence on histological sections for 3 distinctive developmental stages: P4, P10 and P16. *Ncoa3* expression was detected in the cochlea and the organ of Corti with *stria vascularis* in all the time-points (Figure 2A and B). In addition to these structures, immunofluorescence showed cytoplasmic localisation of *Ncoa3* in the Reissner membrane, basilar membrane, spiral limbus and spiral ganglion (Figure 2B). Zebrafish have only one ortholog of *NCOA3*. Whole mount *in situ* hybridization in zebrafish showed *ncoa3* expression in the otic vesicle of 3 and 5dpf zebrafish larvae (Figure 2C). Interestingly, there was continued expression even after the ear system is completely developed, as detected in the inner ear of juvenile fish (5 and 7wpf, weeks post fertilization) (Figure 2C). *ncoa3* expression was not detected in neuromasts (mechanosensory system able to detect small water vibrations). Therefore, our results suggest a conserved expression pattern of *Ncoa3* in the ear.

**Figure 2.**
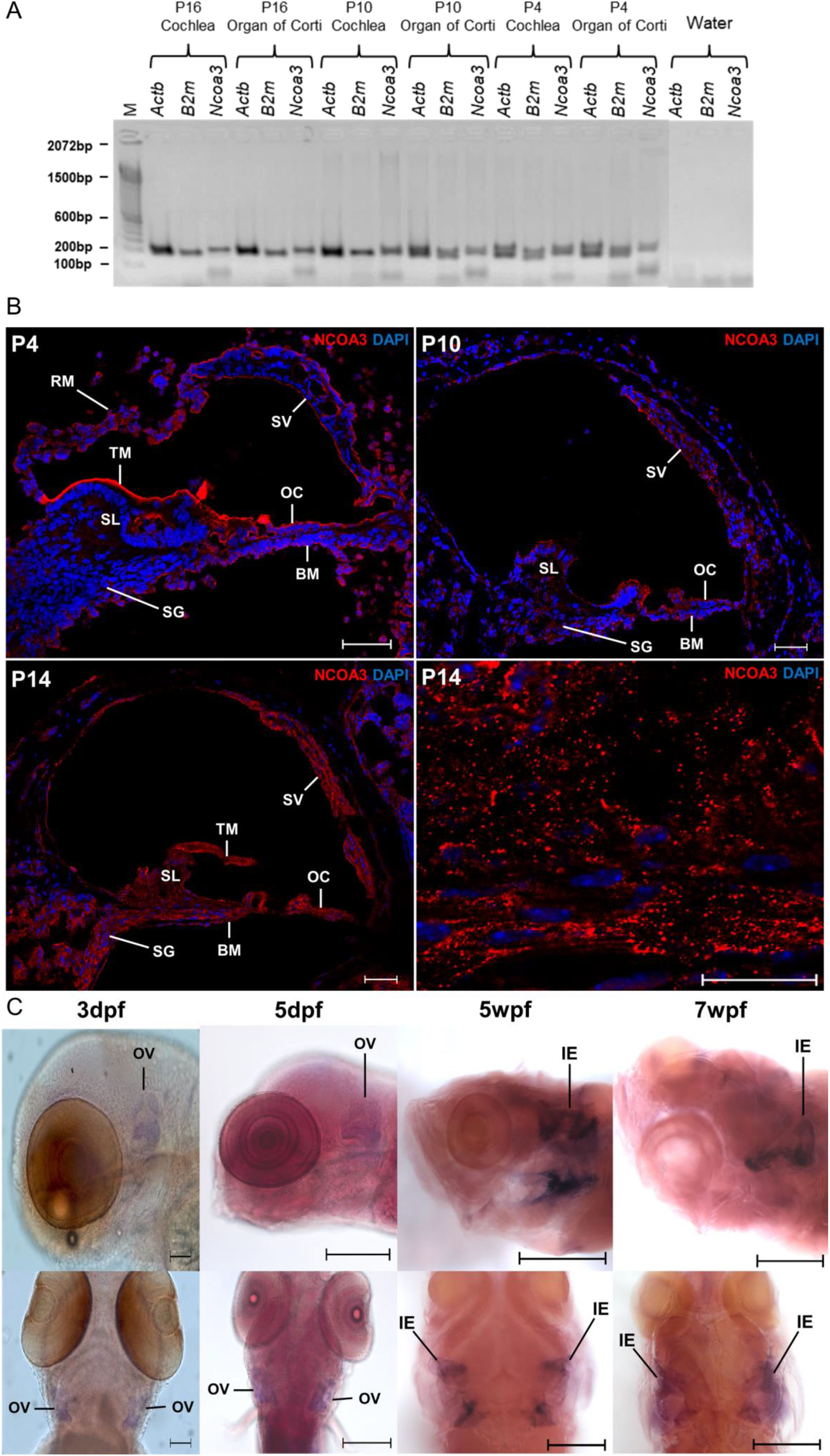
*Ncoa3* is expressed in mice ear at P4, P10 and P16. A) RT-PCR shows expression of *Ncoa3* and housekeeping genes *Actb* and *B2m* for the different stages of mice cochlea development and Organ of Corti. M= 100bp molecular weight. Note that *Ncoa3* is expressed in all stages analysed and in both tissue samples. B) Immunofluorescence on transversal histological sections of mice cochlea. In the bottom right corner, a greater zoom of P14 mice cochlea is displayed, showing expression pattern of *NCOA3.* Anti-NCOA3 (red) has been used, with nuclei shown in blue (DAPI). BM= Basilar Membrane, OC= Organ of Corti, RM= Reissner Membrane, SG= Spiral Ganglion, SL= Spiral Limbus, SV= Stria Vascularis, TM= Tectorial Membrane. C) Expression of endogenous *ncoa3* in zebrafish inner ears at larval stages: 3 dpf and 5dpf (days post-fertilization); and juvenile stages: 5wpf and 7wpf (weeks-post-fertilization). Scale bars= 200μm for 3 and 5 dpf, and 500μm for 5 and 7wpf.

### Zebrafish *ncoa3*^*bi456/bi456*^ show cartilage cell behaviour abnormality in the otic vesicle

In order to investigate the potential role of *NCOA3* in the pathogenesis of hearing impairment, we generated *ncoa3* homozygous zebrafish mutants using CRISPR/Cas9 genome editing. *ncoa3*^*bi456/bi456*^ (*ncoa3−/−)* carry a 5bp deletion (delTACGA) leading to a premature stop codon at position 518aa (S518_Y1520del), reducing the protein size from 1520aa to 517aa. Human and zebrafish sequence alignment showed conservation of 2 out of 5bp within the deletion site. A deleterious effect was predicted when simulating the same mutation in the human ortholog.

*NCOA3* has been previously associated, through GWAS studies, with osteoarthritis, bone mass, abnormal cartilage behaviour, and notch signalling pathway (50–54). To investigate chondrocyte behaviour and sensory cells expressing notch in the zebrafish ear, we crossed *ncoa3*^*bi456/bi456*^ to a double transgenic line carrying Tg(*col2:mcherry; notch:egfp*). Zebrafish *ncoa3*^*bi456/bi456*^ did not display any major morphological abnormalities of the ear at 5dpf (Supplementary Figure 3A and B). Surprisingly, we detected abnormal clusters of cartilage cells (*mcherry* positive) lining the cristae region in 95% of larvae (Figure 3A). 3D image analysis showed tight association of abnormal cartilage cells with notch positive sensory cells (Figure 3A). To examine if detachment of cartilage cells from main cartilage elements (exostosis) was disrupting the hair cells, we measured the lengths of the stereocilia and cupula of the lateral and anterior cristae at 5dpf and no significant differences were detected (Supplementary Figure 3C).

**Figure 3.**
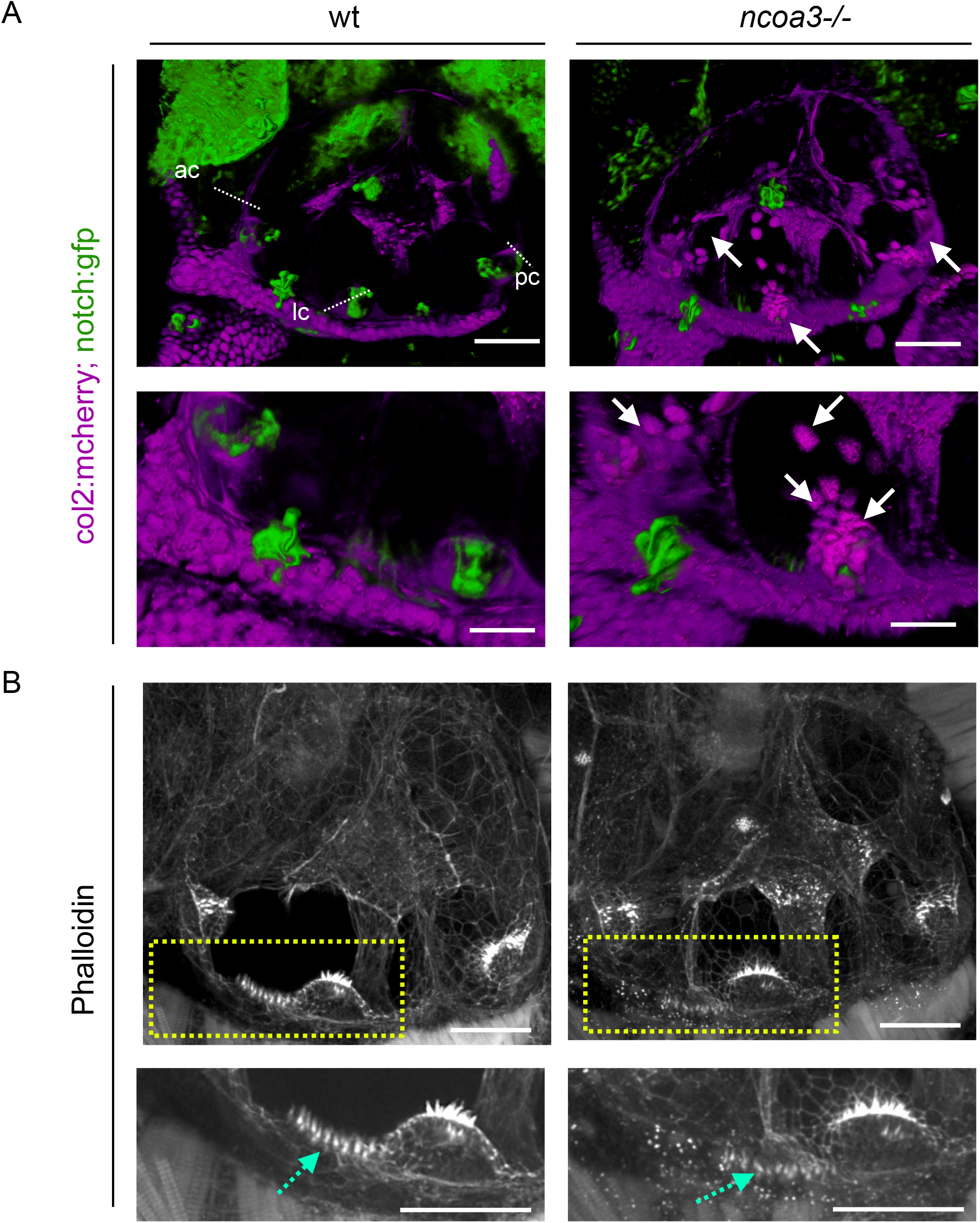
Abnormal cartilage behaviour and macula hair distribution in ears of *ncoa3*−/−. A) 3D renders from confocal images of wt and *ncoa3*−/− carrying Tg(*col2:mcherry, notch:gfp*) to show cartilage and cristae, respectively. Arrows indicate abnormal cartilage cell behaviour (cell exostosis). (ac= anterior crista, lc= lateral crista, pc= posterior crista). Scale bars= 50μm. Regions of anterior crista and macula were zoomed in. Scale bars= 20μm. B) Phalloidin staining and confocal imaging to show the distribution of hair cells. Yellow dashed box to show zoomed in region. Abnormal distribution of hair cells was observed in the macula (dashed cyan arrows). Scale bars= 50μm, zoomed in region = 20μm.

In addition, phalloidin staining, which labels actin filaments of the stereocilia, was performed to evaluate stereocilia of other regions of the ear. Interestingly, we detected disorganised distribution of stereocilia of the macula (4/4 of *ncoa3−/−* and 0/3 wt) (Figure 3B). We investigated earlier stages of development (2-3dpf) to understand when chondrocytes were first misplaced. While at 2dpf no differences were detected, by 3dpf ectopic chondrocytes were observed at the cristae and internal regions of the ear canal (Supplementary Figure 4). This suggests exostosis of cartilage cells from the ear cartilage layer towards the cristae regions and internally. We did not detect changes in the neuromasts throughout the larvae, neither in the lateral line (data not shown). We also did not observe differences in larval swimming behaviour of *ncoa3*^*bi456/bi456*^ at 5dpf (data not shown). Our results suggest abnormal cartilage behaviour (exostosis) and disruption of stereocilia organisation in the macula as a potential progressive and subtle mechanism underlying hearing loss.

### Higher bone density and ectopic mineralisation within the ear of adult *ncoa3*^*bi456/bi456*^

Otoliths consist of a proteinaceous core that is biomineralized by calcium carbonate; in the adult fish ear, a single otolith is tethered to each of the utricular, saccular and lagenal sensory maculae allowing sensation of linear accelerations and sound (55). It has been shown that mutations in *Otogelin* and α-*Tectorin* impair otolith seeding (56), and mutations in their human orthologs *OTOG* and *TECTA* cause deafness. Therefore, the shape and density of otoliths are indicative of possible defects in the hearing system. *ncoa3*^*bi456/bi456*^ survive to adulthood and are fertile. To analyse the 3D structure of the adult ears, we performed micro-computerised tomography (μCT) of 1 year old mutants (n= 8) and wts (n= 25). We observed higher bone mineral density of craniofacial bones of *ncoa3*^*bi456/bi456*^ and abnormal and disorganised mineralisation of amorphous material was detected in 75% (6/8) of the ears of *ncoa3*^*bi456/bi456*^, but was never observed in wt (Figure 4A). This mineralisation was attached to the lagenal otoliths, which is clearly observed through cross sections (Figure 4A, arrows). We did not deteect abnormalities in the utricular and saccular otoliths. Moreover, otoliths showed increased bone mineral density in mutants (Figure 4A and C). Therefore, these results suggest a role of *ncoa3* in bone and ectopic mineralisation regulation in the ears that could lead to progressive hearing impairment in adult fish. It has been shown that vestibular function can be assessed through swimming behaviour analyses (57). Therefore, to test if the fish displayed any signs of hearing loss we analysed swimming behaviour by tracking individual fish in 2D in a tank containting a shaded corner, and calculating the spatial distribution heterogeneity of fish under constant ambient background noise. Vestibular malfunction has been associated to abnormal exploratory behaviour in zebrafish (57). We hypothesised that if hearing function is altered in *ncoa3*^*bi456/bi456*^, these fish would display a distinct exploratory behaviour, dispersing from the shaded corner of the tank more often than the wt. While the wt was retained mostly to the shady corner, interestingly the *ncoa3*^*bi456/bi456*^ showed increased spatial distribution heterogeneity, detected through the comparison of total trajectory distribution between both groups (Figure 5). Our results suggest possible hearing malfunctioning in *ncoa3*^*bi456/bi456*^ due to differences in bone densities, and ectopic mineral deposition.

**Figure 4.**
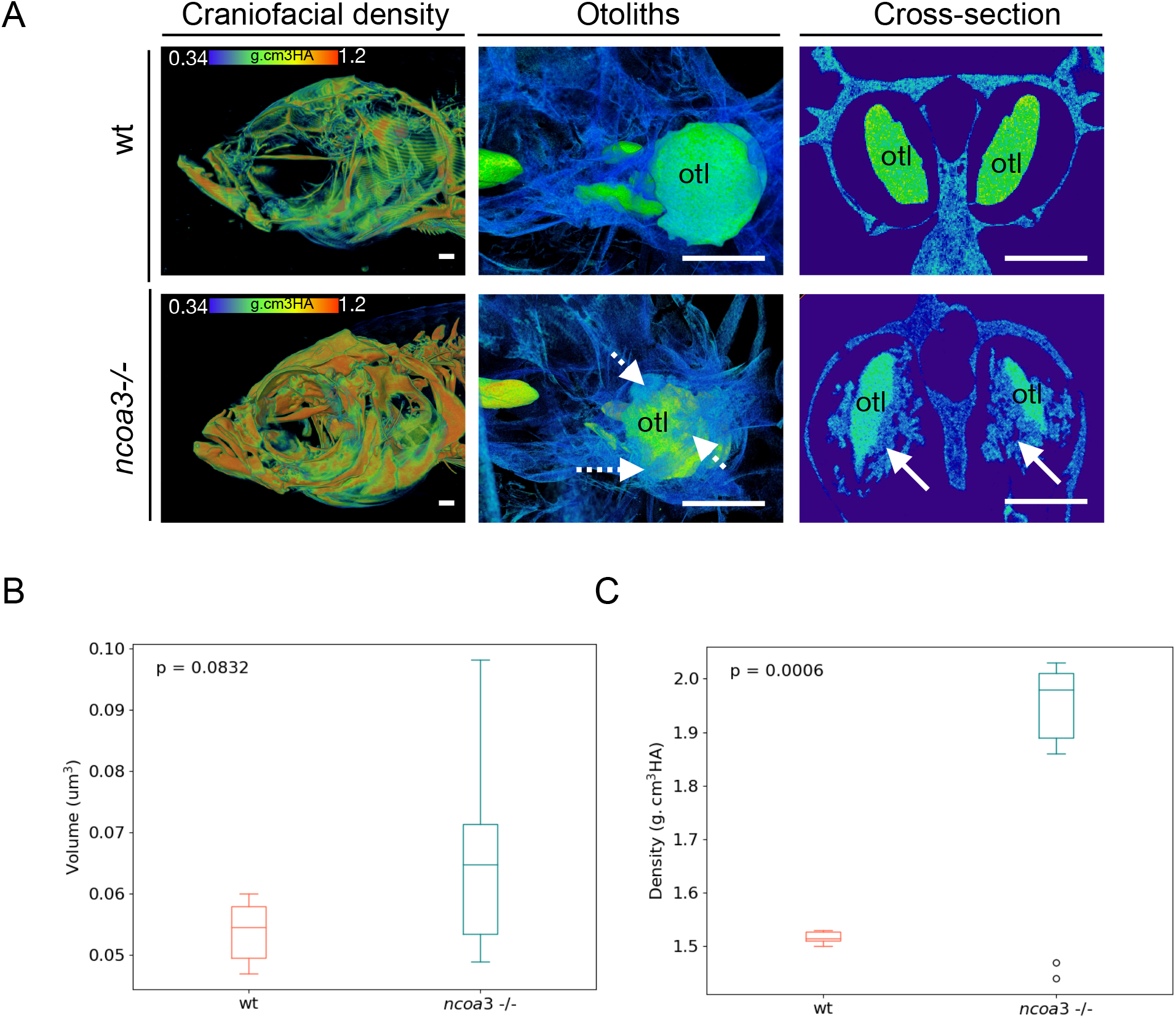
Abnormal mineralisation of amorphous material within the adult inner ears and higher BMD in *ncoa3−/−*. A) 3D renders from μCT images of wt and *ncoa3−/−* of same age (1 year old). The head was color-coded to show bone mineral density (g.cm^3^HA; min= 0.338; max= 1.124). Note that craniofacial bones in *ncoa3* mutants have higher density compared to wt. Otoliths (otl= arrows) were zoomed in. Abnormal mineralisation (dashed arrows) is observed attached to the otoliths. A cross section picture was taken to show the mineralised amorphous material (arrows) juxtaposed to the otoliths. B) Volume of otoliths. C) Bone mineral density of central region of otoliths. Non-parametric, two-tailed, independent Student’s t-Test was used as statistical analysis (p<0.05). Scale bars= 500μm.

**Figure 5.**
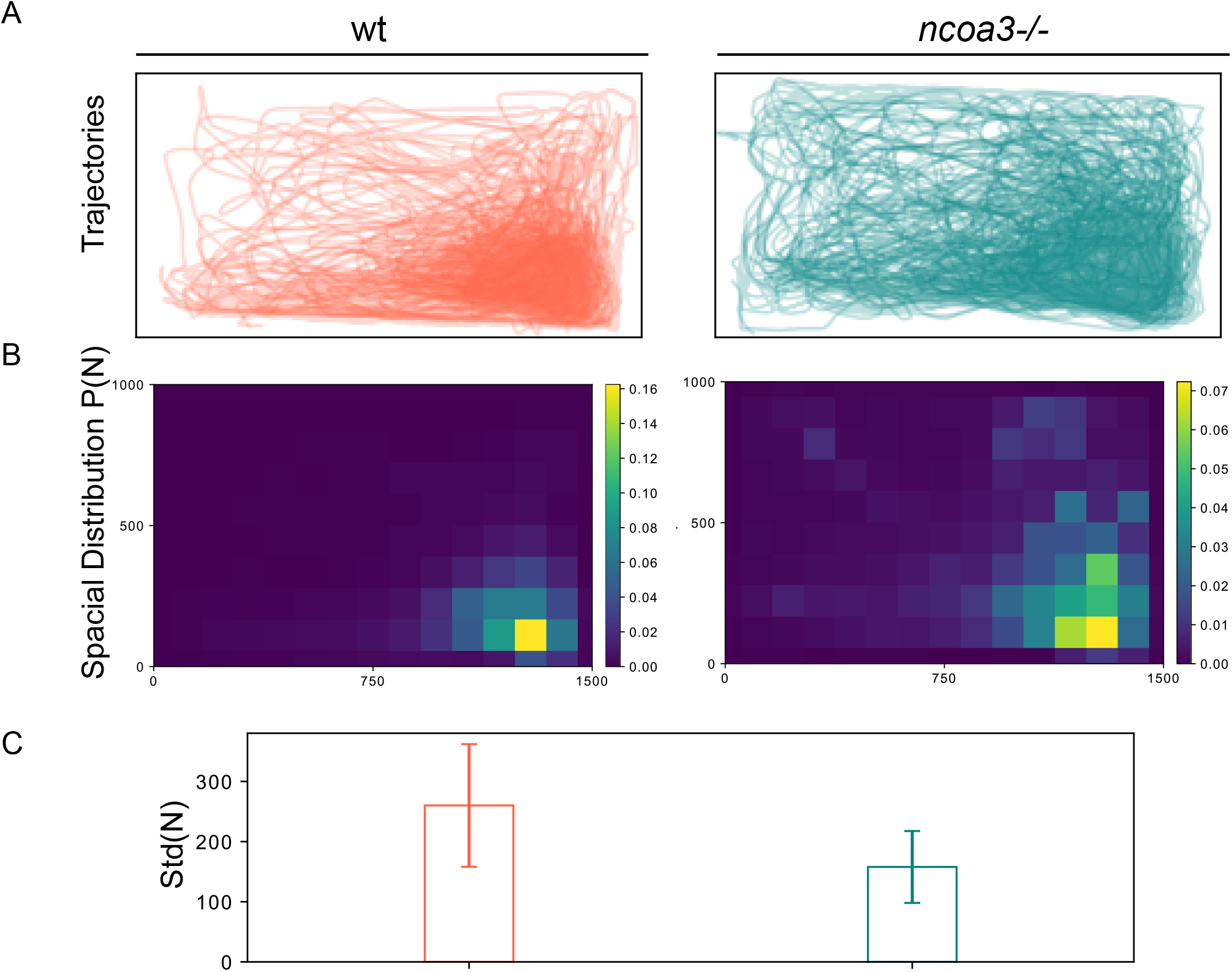
Altered swimming behaviour of adult *ncoa3 −/−* suggests hearing malfunction. A) Overlapped Trajectories of 1-year old wt (n= 7) and *ncoa3−/−* (n= 6). The bottom right corner is shaded, where the fish are more likely to stay. B) Average spatial distribution of wt and *ncoa3−/−*. A brighter colour indicates that the fish are more likely to stay in the respective region. For every trajectory acquired from different fish, the corresponding spatial distribution P(N) was calculated. From every P(N), the standard deviation of N in all the grids is calculated, noted as Std(N). C) Graph of Std(N) values for wt and *ncoa3−/−*. Error bars represent the standard deviation of Std(N). Non-parametric, two tailed, t-Test was used (p =0.06).

## Discussion

Non-syndromic hearing loss is a condition that affects almost 466 million people worldwide and is characterized by a broad heterogeneity of causes, among them genetic factors play an outstanding role (2). Reported genes and variants associated with monogenic forms of the disease have been identified from independent pedigrees and the rarity of some variants implies that some candidates are hardly reproducible. Therefore, functional studies are key to support genetic findings. In this sense, animal models such as mice and zebrafish are valuable and have proved relevant to the investigation of molecular mechanisms that underlie hearing loss in humans (56,58–60). The molecular and cellular mechanisms involved in ear development and homeostasis are highly conserved through evolution and zebrafish have been used elsewhere to study ear development and to confirm candidate genes involved in hearing loss (61–65).

Here, combining linkage analysis with exome sequencing and functional analysis we have reported for the first time an association between segregation of a rare variant in *NCOA3* and hearing loss, suggesting a novel mechanism leading to the pathogenesis of hearing impairment. NCOA3 is a nuclear receptor coactivator from the *NCOA* gene family, positively regulating nuclear receptor-mediated gene transcription (66). We identified a missense variant in *NCOA3*, c.2810C>G: p.Ser937Cys, of which computation predictions and frequency are compatible with the hypothesis of this variant being causative of hearing loss. We have provided further expression data in mice and zebrafish ears, that point to evolutionary conservation of gene function in the ear. Moreover, through CRISPR/Cas9 we have generated a *ncoa3* zebrafish knockout to further investigate the effects of loss of function of *ncoa3* in the ear during development and ageing.

NCOA3 function has been linked to reproductive development and physiology regulation (67–70), pluripotency regulation (71), neurotransmitter metabolism regulation (72), adipogenesis promotion (73), long-chain fatty acid metabolism regulation (47). In mice, although around 10% of the knockout animals for *Ncoa3* exhibit a unilateral drop of the ear (74), they were not submitted to audiological evaluation. In a previous transcriptome analysis study of mice tissue, *Ncoa3* expression has been reported in organ of Corti of E16, P0, P4 and P7 C57BL/6, with more pronounced expression levels observed in the postnatal phases (48). Transcriptome analysis of inner and outer hair cells from P25-P30 CBA/J mice cochleae also indicated expression of *Ncoa3* (49). Our results not only confirmed *Ncoa3* expression in the organ of Corti of P4 mice, but also complemented the aforementioned studies, showing that the gene is still active in more advanced ages, near the end of cochlea maturation (P10 and P16). Moreover, we showed evidence of NCOA3 protein expression in the mice hearing system. Protein expression has been detected in all mice ages studied (P4, P10 and P14), with expression pattern spread along several cochlear structures: basilar membrane, Reissner membrane, organ of Corti, *stria vascularis*, spiral limbus and spiral ganglion. Altogether, our results suggest that *Ncoa3* may have an important role in the development and physiology of mice auditory system.

We also detected expression of *ncoa3* in zebrafish during and after the completion of the inner ear development. In addition to the inner ear, fish have another component to their mechanosensory system; the lateral line, which is also formed by hair cells, supporting cells and sensory neurons, forming units called neuromasts which are key during the startle swim behaviour response (75,76). Although there are genes that are expressed both in the inner ear and lateral line, such as *atoh1a* (77,78) and *ngn1* (79) and that larval behaviour is observed when such genes are knocked out, *ncoa3* does not follow this pattern, as we did not detect its expression in the neuromasts or changes to larval startle swim behaviour response in zebrafish *ncoa3*^*bi456/bi456*^.

Functional analysis carried in zebrafish *ncoa3*^*bi456/bi456*^ showed that *ncoa3* is dispensable for development of the inner ear, but it is important for the maintenance the skeletal system. Although we did not detect morphological changes (size and shape of ear) in larval stages, abnormal cartilage cell behaviour was a predominant phenotype in larval *ncoa3*^*bi456/bi456*^. In adults, denser craniofacial bones and otoliths, and ectopic mineralisation in the ears were detected. Abnormal invading cartilage cells could potentially contribute to ectopic mineralisation in the vestibular region during ageing. Recent studies inferred *NCOA3* involvement in maintaining skeletal homeostasis, with evidence of its function in bone mass (53), behaviour and molecular signature of chondrocytes (80,81).Thus, sustaining its role in regulation of bone density and cartilage behaviour, respectively. Changes in bone mineral density have also been associated with hearing loss. Loss of bone mineral density in the cochlea capsule has been related to hearing loss in Paget’s disease (osteoclast/bone resorption disorder (82). Mutations in *SOST* (sclerostosis and van Buchem’s disease) cause enhanced bone formation, higher bone mineral density, and calvaria overgrowth, which frequently compresses cranial nerves leading to hearing loss (83). Although computed tomography scans of temporal bones revealed normal bone morphology in affected individuals from the pedigree, it would be interesting to further investigate overall calvarial bone thickness and bone mineral density in the family, as such data are currently unavailable. Moreover, computed tomography is not sensitive enough to detect possible subtle changes at the cellular level that could be contributing to hearing loss as suggested by our functional analysis.

Altered swimming behaviour was previously detected when mutant larvae for several hearing loss associated genes were analysed, such as *grhl2b*, *myo7aa*, *cdh23*, *otofa* and *otofb* (84,85). Mutations in the human orthologs are associated with mild to severe hearing loss (86–89). The respective zebrafish mutants have severe abnormalities in the inner ear, otoliths and/or lateral line, and recapitulate abnormalities of those observed in human patients. However, they differ from subtle and progressive changes involved in *ncoa3* zebrafish mutants and the family that we described. We did not observe larval behaviour changes in *ncoa3*−/− (data not shown). But we observed adult behaviour changes that fit with progressive hearing loss. While assessment of hearing loss through adult swimming behaviour in zebrafish is not well explored yet, it has been shown that when adult zebrafish are introduced into a centre of a magnetic field they exhibited altered exploratory behaviour due to vestibular malfunction and independent of lateral line function (57). Therefore, vestibular function can be assessed by exploratory behaviour changes. In a new environment under constant background noise, we would expect that fish carrying hearing disability would display altered behaviour. Ectopic mineral deposition within the ears of adult mutant zebrafish and increased density of otoliths and craniofacial bones are potentially correlated with the altered vestibular function and swimming behaviour found in adult mutants.

Although family size does not allow a definite conclusion about the c.2810C>G variant being causative to hearing loss, our functional results were compatible with the hypothesis of *NCOA3* playing a role in hearing, suggesting skeletal homeostasis (cartilage behaviour and bone density) as a strong factor involved in the condition. Our contribution was to attract further attention to *NCOA3* as possibly involved in hearing, since many groups are dealing with patient samples revealing hundreds of candidate variants after exome sequencing, without clues to find the causative one. Further functional studies to evaluate the precise effect of the missense variant p.Ser937Cys in *NCOA3* function would add value in understanding age-related hearing loss in patients with autosomal dominant pathogenic variants in *NCOA3*.

## Materials and Methods

### Patients

A large Brazilian family comprising 5 generations and 15 reported affected individuals with hearing loss was ascertained in our genetic counselling unit (Centro de Pesquisas sobre o Genoma Humano e Células-Tronco - IBUSP) for molecular studies. The transmission of hearing loss in the pedigree is compatible with autosomal dominant inheritance (Figure 1A). For molecular studies, DNA samples from 19 individuals were collected: 7 from affected individuals (III-5, III-8, III-10, IV-6, IV-9, IV-21, IV-22), and 12 from unaffected individuals, including spouses (II-4, III-11, IV-5, IV-12, IV-15, IV-17, IV-18, IV-19, IV-20, V-3, V-5 e V-6). Written informed consent was obtained from every participant or the respective guardians. The study was approved by the Ethics Committee from Instituto de Biociências da Universidade de São Paulo.

### Audiological evaluation

Pure tone audiometry, both air (frequencies ranging from 250 to 8000Hz) and bone conduction (frequencies ranging from 500 to 4000Hz) were performed for identification of hearing threshold levels in seven affected individuals (III-5, III-8, III-10, IV-6, IV-9, IV-21 and IV-22) and eleven non-affected individuals (III-11, IV-5, IV-12, IV-15, IV-17, IV-18, IV-19, IV-20, V-3, V-5 and V-6). Most of these exams were done at DERDIC (Divisão de Educação e Reabilitação dos Distúrbios da Comunicação, PUCSP), while some were conducted by other institutions prior to this study.

### SNP-array and microsatellite markers genotyping

Genomic DNAs from seven affected individuals (III-5; III-8 III-10; IV-6; IV-9; IV-21; IV-22) were submitted to SNP-Array (50K) assays (Affymetrix GeneChip HumanMapping 50K Array, Affymetrix), using the manufacturer’s reagents (XbaI) and following the GeneChip Mapping 10K 2.0 Assay Manual. Scanning was performed in a Genechip Scanner 3000 and interpreted with Affymetrix Genotyping Console software (Affymetrix). In addition, twelve polymorphic microsatellite markers mapped to chromosome 20 (ABI Prism Linkage Mapping Sets v2.5) were genotyped in 16 samples (II-4, III-5, III-8, III-10, IV-5, IV-6, IV-9, IV-12, IV-16, IV-17, IV-18, IV-19, IV-20, IV-21, IV-22, V-3).

### Lod score calculations

Penetrance of hearing loss was estimated according to methods previously described (19). The most likely value of penetrance was K= 0.6364. Multipoint logarithm of odds (LOD) score values were calculated, for each autosome, using Merlin program (20) under dominant inheritance model, assuming a rare allele (frequency= 0.001). The LOD score calculations were performed considering penetrance of K= 0.6364, but also under the assumption of penetrance K= 0.8 and complete penetrance, K= 1.

### Whole-exome sequencing

DNA samples from two affected individuals (III-8 and III-10) were submitted to whole-exome sequencing. The library was prepared with Nextera rapid capture kit (Illumina), sequence capture was performed with Illumina Exome enrichment kit (~62 Mb target size) and sequencing was performed using HiSeq 2500. Fastq files were aligned against reference GRCh37 with Burrows-Wheeler Aligner (BWA) (21), realignment of indel regions, discovery of variants and recalibration of base qualities were performed using GATK software (22) for the production of VCF files; the VCF was annotated by ANNOVAR software (23). Variant frequencies were compared with public variant databases: 1000 Genomes (24), National Heart, Lung, and Blood Institute Exome Sequencing Project (NHLBI-ESP) (25), Genome Aggregation Database (gnomAD) (26) and Online Archive of Brazilian Mutations (ABraOM) (27). Polyphen-2 (28), SIFT (29), Provean (30) and MutationTaster2 (31) were used for *in silico* damage prediction to the protein. Protein sequence alignment near the best candidate variant was performed by Clustal Omega alignment program (32).

### Sanger sequencing

The DNA regions containing candidate variants filtered after exome sequencing were amplified by PCR. The products were bi-directionally Sanger sequenced with the BigDye Terminator v3.1 Cycle Sequencing Kit (ThermoFisher Scientific) in ABI 3730 DNA Analyzer (Applied Biosystems). *NCOA3*F-5’GGCTGTACTTACATGGTATAAGAAGG3’, *NCOA3*R-5’AGGGGAGGGTGGACACTTAC3’, *DIAPH3*F-5’CAAGGGTTTCTGTGCATACC3’, *DIAPH3*R - 5’CACTACTCGTTAGTAAATGGAAGGG3’, 5’GCTGAGAAGACCACCTGCAT3’, *USH1C*F - *USH1C*R-5’GAGGAGGAGGAAGTTGGCTG3’ were used as primers. Sequences were analysed using Bioedit (Ibis Biosciences).

### Multiple alignment of *NCOA3* and its orthologous

Multiple alignment of *NCOA3* gene and protein with its orthologous was performed using Clustal Omega provided by European Bioinformatics Institute (EMBL-EBI) (33). For this purpose, the following sequences were used: *Homo sapiens* (NM_181659.2 and NP_858045.1); *Pan troglodytes* (XM_016938072.2 and XP_016793561.2); *Macaca mulatta* (XM_015148801.1 and XP_015004287.1); *Bos Taurus* (XM_002692493.4 andXP_002692539.1;*Mus musculus* (NM_008679.3 and NP_032705.2); *Rattus norvegicus* (XM_006235634.2 and XP_006235696.2); *Gallus gallus* (XM_004947056.2 and XP_004947113.2); *Danio rerio* (XM_687846.9 and XP_692938.5) *Xenopus tropicalis* (XM_018097860.1 and XP_017953349.1).

### Mice husbandry

CBL57/6 mice were obtained from Centro de Pesquisas sobre o Genoma Humano e Células-Tronco (IBUSP) experimentation housing facility. The animals were housed as previously described by Council for International Organizations of Medical Sciences (CIOMS) (34). All experiments with mice were ethically approved by the Internal Review Board on Ethics in Animal Research from the Instituto de Biociências da Universidade de São Paulo (Process Number 16.1.668.41.6).

### Cochleae and organ of Corti dissection

Cochleae and organ of Corti with *stria vascularis* were harvested from CBL57/6 decapitated mice at 4, 10, 14 and 16 day-old (P4-P16) postnatal CBL57/6 mice. After decapitation, the head was bathed in ethanol 70%, followed by longitudinal incision at the skull’s sagittal line and visualization of the temporal bone, allowing the dissection of the labyrinth. For RNA extraction, the labyrinths were transferred to a Petri dish with RNAlater® (Sigma Aldrich). Cochlea and organ of Corti with *stria vascularis* were then surgically harvested with micro tweezers (Dumont #5 e #54, Koch Electron Microscopy) under trinocular stereomicroscope (Discovery V12, Carl Zeiss). For immunofluorescence assays, cochleae were isolated from labyrinths kept in phosphate buffered saline (PBS), using micro tweezers (Dumont #5 e #54, Koch Electron Microscopy) under trinocular stereomicroscope (Discovery V12).

### RT-PCR

Total RNA extraction was performed with a pool of 12 cochleae or 12 organs of Corti with *stria vascularis* from P4, P10 and P16 mice, as well as with gastrocnemius sample of P180 mice. Total RNA was extracted with RNeasy Microarray Tissue Mini Kit (QIAGEN). Synthesis of cDNA was performed with RNeasy Microarray Tissue Mini Kit (QIAGEN), using 1μg of total RNA. Primers used for this experiment were: *Ncoa3*F-5’CGTTTCTCCTTGGCTGATGG3’, *Ncoa3*R-5’CGGGATTTGGGTTTGGTCTG3’, *Actb*F-5’GGCTGTATTCCCCTCCATCG3’, *Actb*R-5’CCAGTTGGTAACAATGCCATGT3’, *B2m*F-5’TCGCGGTCGCTTCAGTCGTC3’, *B2m*R-5’TTCTCCGGTGGGTGGCGTGA3’. Control experiments concomitantly performed were negative control of cDNA synthesis (using water instead of extracted RNA), negative control of RT-PCR (using water instead of cDNA), and positive control (using cDNA synthetized from gastrocnemius RNA). Housekeeping genes used as reference for this experiment were *Actb* and *B2m*.

### Immunofluorescence assays

Cochleae preparation and immunofluorescence assays were performed as described by (35). Cochleae were perfused locally and fixed in 4% paraformaldehyde (PFA) at 4°C overnight (o/n). P10 and P14 passed through decalcification with 10% EDTA and 1% PFA at 4°C for 4 days. All tissues were washed with 1X PBS, submitted to serial dilutions of sucrose solution and Jung Tissue Freezing Medium (Leica Microsystems), frozen and transversely cryosectioned in 12μm. Slides were stored at - 80°C until use. For immunofluorescence assays, histological slides were simultaneously permeabilized and blocked with 0.3% triton X-100 and 4% bovine serum albumin (BSA) solution, followed by incubation in solution containing 1:50 polyclonal anti-NCOA3 antibody (anti-SCR3 antibody - ChIP Grade, Rabbit Polyclonal, ab2831, Abcam Inc.) diluted in 0.1% triton X-100 and 4% BSA, at 4°C o/n. Subsequently, the slides were incubated in solution containing 1:500 anti-rabbit AlexaFluor-568 diluted in 0.1% triton X-100, 1% BSA, at for 2h at room temperature. After rinse in PBS, the slides were then mounted with Prolong Gold Antifade Reagent (Invitrogen) with DAPI for nuclei staining. Images were taken confocal microscope either LSM 780 (Carl Zeiss) or LSM880 (Carl Zeiss), using Zen software (Carl Zeiss).

### Zebrafish husbandry and lines

Zebrafish were housed as previously described (36). Animal experiments were ethically approved by the Animal Welfare and Ethical Review Body (AWERB) at the University of Bristol and performed under a UK Home Office project and by the Internal Review Board on Ethics in Animal Research from the Instituto de Biociências da Universidade de São Paulo (Process Number 16.1.668.41.6). Transgenic lines used have been previously described: *TgBAC(Col2a1a:mCherry)*^*hu5910*^(37) and *Tg(notch:egfp)* (38).

### Whole-mount *in situ* hybridization in zebrafish

Whole-mount *in situ* hybridizations on zebrafish samples were performed as described by (39). *ncoa3 in situ* probe was synthesised *in vitro* from a PCR product(880 bp amplified from exon11) using a T7 RNA Polymerase for transcription (ThermoFisher Scientific) and DIG-labelling Mix (Roche)followed by purification with SigmaSpin™ Post-Reaction Clean-Up Columns (Sigma Aldrich). The following primers were used for the PCR: *ncoa3-F* (5’GAATACCTTCTCTAGCAGCTCATTG3’) and *ncoa3-R* (5’taatacgactcactatagggagCTTATTGAGGAGGTAGTGAAGGAGG3’).

### Generation of zebrafish *ncoa3−/−*

*ncoa3* mutants were generated by CRISPR/Cas9 system as previously described (40). A gRNA was designed to target *ncoa3:* TGGGGTCTCCGCGGATACGA<u>GGG</u>(PAM) (chr11:18516059-18516081). Once synthesised, it was incubated with GeneArt Platinum Cas9 nuclease (Invitrogen) prior to injections into 1 cell stage zebrafish embryos. DNA was extracted from 20 individual larvae at 2 days post fertilization (dpf), followed by PCR amplification (*ncoa3*CRISPR F: FAM-ATGAATGAGCAAGGCCACAT; *ncoa3*CRIPSR R: GGACTTGCTCCCATTTTAGG) and subjected to fragment length analysis (ABI 3500) to test gRNA efficiency (90% efficiency rate detected). G0s were outcrossed to generate G1s which were submitted to Sanger sequencing. The mutant line *ncoa3*^*bi456*^ carries a 5bp deletion, leading to a premature stop codon predicted to undergo mRNA nonsense mediated decay.

### Microscopy

Samples were mounted in 1% low melting point (LMP) agarose (Invitrogen) and imaged with a Leica SP5II confocal microscopy (Leica LAS software) using 10x PL APO CS (dry), 20x immersion lens (phalloidin) or 40x PL APO CS (oil) lenses (cristae imaging). LasX (Leica) and Amira 6.0 (FEI) was used for 2D and 3D rendering, image analysis and picture acquisition.

### 3D perspective measurements of the otic vesicle

Two distinct 3D perspective measurements (sagittal/x axis and coronal/y axis) were taken of the major axis of the otic vesicle at 5dpf from confocal images using Amira 6.0. GraphPad (Prism) was used for statistical analysis. t-Tests (non-parametric, Mann-Whitney U test, p< 0.05) were performed (n= 7 for each group).

### Phalloidin staining

Larvae (5dpf) were fixed in 4% PFA at 4°C o/n, washed in PBS 3 × 5 minutes and incubated in AlexaFluor 555 conjugated phalloidin (1:20 in PBS) o/n at 4°C (protocol adapted from (41)). Samples were then washed in PBS 3 × 15 minutes, mounted laterally in 0.5% LMP agarose and imaged on a confocal microscopy.

### Micro-computerised tomography (μCT) and bone mineral density calculation

Adult fish (1 year old) were fixed in 4% PFA for 1 week followed by sequential dehydration to 70% ethanol. Fish heads were scanned using an XT H 225ST μCT scanner (Nikon) with voxel size of 20μm and 5μm for detailed geometric analysis, using an x-ray source of 130 kV, 53μA and without additional filters. Images were reconstructed using CT Pro 3D software (Nikon). Amira 6.0 was used for image analysis and to generate 3D volume and surface renders. Lagenal otoliths were segmented from μCT images (20μm resolution) and bone density was quantified by sampling a region 20 slices thick at the centre of the otoliths. Greyscale values were calibrated against phantom blocks with calibrated densities.

### Adult swimming behaviour analysis

Swimming behaviour was assessed by recording 2D movement (from above) in a rectangular tank. Individual fish (1 year old, wild type (wt) n= 7 and *ncoa3−/−* n= 6) were transferred to a tank (35cm × 40cm) containing a shaded corner (10cm × 10cm) and a total of 8 L of water. The tank was placed in an environment of constant background noise (~80dB) and recorded with a digital camera (Balser acA2040-120μm) at a frame rate of 15 frames/s, for 10 minutes (9000 frames). The 2D positions in different frames were calculated by a custom Python script (42). Trajectories were obtained by applying a four-frame best estimate algorithm (43) to the positions, and further refined following the approach described in (44). For each fish, we calculated its normalised 2D spatial distribution P(N) where N is the number of times the fish stays in a grid, whose size is 100 pixels by 100 pixels. We then calculated the standard deviation of N in all grids as a proxy to the spatial distribution heterogeneity, noted as Std(N). The difference in the behaviours between the wt fish and the mutant fish is then characterised by Std(N) values of each individual. Assuming the spatial heterogeneity of the wt fish and the mutant fish follow the normal distributions with different variances, we used student t test to calculate the probability of their average values being the same (p< 0.05 statistically significant).

## Acknowledgements

EK and CH were funded by Versus Arthritis (19476, 21211). AK was funded by BBSRC SWBio-DTP studentship and EL by a Wellcome Trust Dynamic Molecular Cell Biology PhD studentship. This work was supported by Fundação de Amparo à Pesquisa do Estado de São Paulo (FAPESP - CEPID Human Genome and Stem-Cell Research Center 2013/08028-1), Coordenação de Aperfeiçoamento de Pessoal de Nível Superior - CAPES and Conselho Nacional de Desenvolvimento Científico e Tecnológico (CNPq, grant 133182/2015-0). The authors are indebted to all professionals of Divisão de Educação e Reabilitação dos Distúrbios da Comunicação (DERDIC) da Pontifícia Universidade Católica de São Paulo, São Paulo (PUC-SP), in special to Márcia Zucheratto, for audiological evaluations. We thank Dr. Maria Rita Passos-Bueno for support on establishing the zebrafish laboratory at Instituto de Biociências da Universidade da São Paulo (IBUSP) and Dr. Marília O. Scliar for bioinformatics assistance. We also thank Ms. André S. Bueno for assistance and Drs. Erika Freitas and Carla Rosenberg for Array-CGH.

## Author contributions

RSS, VLGD, FSN, BCN, EL, AK, YY and EK performed experiments. RSS, VLGD, LUA, ACB, JO, EL, AK, YY, RCMN and EK analysed data. The project was designed by RCMN (gene identification) and EK (functional analysis). All authors contributed to drafting the manuscript.

## Abbreviations

AWERB: Animal Welfare and Ethical Review Body
ADNSHL: Autosomal dominant non-syndromic hearing loss
BSA: Bovine serum albumin
CIOMS: Council for International Organizations of Medical Sciences
CRISPR: Clustered regularly interspaced short palindromic repeats
Cas9: CRISPR associated protein 9
dpf: Days post fertilization
ENT: Ear, nose and throat
EMBL-EBI: European Bioinformatics Institute
gnomAD: Genome Aggregation Database
LOD: Multipoint logarithm of odds
NHLBI-ESP: National Heart, Lung, and Blood Institute Exome Sequencing Project
ABraOM: Online Archive of Brazilian Mutations
PFA: Paraformaldehyde
P: Postnatal Day
Wpf: weeks post fertilization

