## SUPPLEMENTAL for "*NCOA3* identified as a new candidate to explain autosomal dominant progressive hearing loss"

**Supplementary Materials**

**Supplementary Figure 1. Microsatellite markers flanking *NCOA3*.** Pedigree showing the segregation of *NCOA3* variant (NM_181659: c.2810C>G: p.Ser937Cys) and flanking microsatellite markers haplotype.

**Supplementary Figure 2. Exonic variants found in *DIAPH3* and *USH1C* do not segregate with hearing loss in the family.** Pedigree showing segregation pattern of NM_001258370: c.A1565G:p.Gln522Arg (*DIAPH3*) and NM_005709: c.G946C:p.Glu316Gln (USH1C).

### Supplementary Figure 3. Major morphological abnormalities are not detected in ears of *ncoa3-/-* at 5dpf. A) 5dpf confocal image (maximum projection) of Tg(*col2:mcherry, notch:gfp*). Measurements of sagittal and coronal distances of the otic vesicle were taken at the major axis of the cartilage element (y and x axis). Scale bar = 50 µm. A diagram showing the measurements and position of cristae in the otic vesicle is presented (ac= anterior crista, lc= lateral crista, pc= posterior crista). B) Statistical differences were not detected between wt and *ncoa3-/-* ear morphology. T-tests (nonparametric) (p< 0.05) was used. C) 5dpf confocal images (40x lenses, transmitted light TL) of *ncoa3* mutant (n= 5) and wt (n= 3) carrying Tg(*col2:mcherry*). The lengths (dashed line) of stereocilia (sc) and cupula (c) were taken for the lateral and anterior cristae (images showing lateral cristae). Scale bars = 20µm. D) *ncoa3*-/- did not show statistical differences in lengths of stereocilia or cupula of the lateral and anterior cristae. Multiple t-tests (false discovery rate=1%).

### Supplementary Figure 4. Abnormal cartilage behaviour is detected early in ear development of *ncoa3*-/-. Confocal images of wt and *ncoa3-/-* carrying Tg(*col2:mcherry;notch:gfp*) were taken at 3dpf (n= 5 for each group). Arrows pointing to abnormal cells. Note that abnormal cell migration towards the middle region of the ear is not yet seen at 3dpf. Scale bars= 50µm.

**Supplementary Table 1** – 162 Exome variants found in common in two affected family members, after initial filtration steps described in Table1. B= Benign, D= Damaging, N= Neutral, NA= Not Available, P= Pathogenic, T= Tolerable.

**Supplementary Tables**

Supplementary Table 1 – 162 Exome variants found in common in two affected family members, after initial filtration steps described in Table1. B= Benign, D= Damaging, N= Neutral, NA= Not Available, P= Pathogenic, T= Tolerable.

|  |  |  | **Frequency** | | | **Predicted pathogenicity** | | |
| --- | --- | --- | --- | --- | --- | --- | --- | --- |
| **Chromosome** | **Variant** | **dbSNP** | **ESP6500** | **1kgenome** | **gnomAD** | **SIFT** | **Polyphen2** | **Mutation Taster** |
| chr1 | AGRN:NM_198576:exon16:c.G2737A:p.V913M | rs140120617 | 0,0300% | 0,0000% | 0,0200% | D | D | D |
| chr1 | AMIGO1:NM_020703:exon2:c.G1355A:p.G452D | rs150743133 | 0,6100% | 0,1398% | 0,3900% | D | B | D |
| chr1 | DCDC2B:NM_001099434:exon8:c.C879A:p.H293Q | rs11590012 | 0,7100% | 0,5791% | 0,8100% | T | P | N |
| chr1 | DENND2C:NM_198459:exon2:c.G380A:p.C127Y,DENND2C:NM_001256404:exon4:c.G380A:p.C127Y | rs143842563 | 0,4500% | 0,2995% | 0,0900% | D | B | N |
| chr1 | DENND2D:NM_001271833:exon7:c.C692T:p.S231F,DENND2D:NM_024901:exon7:c.C701T:p.S234F | rs150029631 | 0,4100% | 0,5990% | 0,1100% | T | P | N |
| chr1 | EXO1:NM_003686:exon7:c.G820A:p.G274R,EXO1:NM_006027:exon7:c.G820A:p.G274R,EXO1:NM_001319224:exon8:c.G820A:p.G274R,EXO1:NM_130398:exon9:c.G820A:p.G274R | rs149397534 | 0,2700% | 0,0998% | 0,2100% | D | D | D |
| chr1 | HCN3:NM_020897:exon5:c.G1192T:p.D398Y | rs200186877 | 0,0200% | 0,0000% | 0,0041% | D | P | D |
| chr1 | KCNC4:NM_001039574:exon2:c.G1493A:p.R498Q,KCNC4:NM_004978:exon2:c.G1493A:p.R498Q | rs376704379 | 0,0077% | 0,0000% | 0,0028% | T | B | D |
| chr1 | LRRC8C:NM_032270:exon3:c.A1714G:p.I572V | rs139429089 | 0,0400% | 0,5192% | 0,8400% | T | B | D |
| chr1 | PHTF1:NM_001323049:exon5:c.A373G:p.I125V,PHTF1:NM_006608:exon5:c.A373G:p.I125V,PHTF1:NM_001323041:exon6:c.A373G:p.I125V,PHTF1:NM_001323042:exon6:c.A373G:p.I125V,PHTF1:NM_001323043:exon6:c.A373G:p.I125V,PHTF1:NM_001323044:exon6:c.A247G:p.I83V,PHTF1:NM_001323047:exon6:c.A373G:p.I125V,PHTF1:NM_001323048:exon6:c.A373G:p.I125V,PHTF1:NM_001323051:exon6:c.A247G:p.I83V,PHTF1:NM_001323052:exon6:c.A373G:p.I125V,PHTF1:NM_001323053:exon6:c.A373G:p.I125V,PHTF1:NM_001323045:exon7:c.A247G:p.I83V,PHTF1:NM_001323050:exon7:c.A247G:p.I83V | rs35311195 | 0,4400% | 0,4992% | 0,1200% | T | B | N |
| chr1 | S100A7A:NM_176823:exon2:c.C101A:p.T34K | rs377370756 | 0,0077% | 0,0000% | 0,0012% | T | B | N |
| chr1 | SGIP1:NM_001308203:exon18:c.G1420A:p.V474I,SGIP1:NM_001350218:exon19:c.G1438A:p.V480I,SGIP1:NM_001350217:exon21:c.G2023A:p.V675I,SGIP1:NM_032291:exon21:c.G2011A:p.V671I | rs140384550 | 0,0800% | 0,0599% | 0,0800% | T | P | D |
| chr1 | SYNC:NM_001161708:exon2:c.C787T:p.R263C,SYNC:NM_030786:exon2:c.C787T:p.R263C | rs41265855 | 0,2200% | 0,2596% | 0,3700% | D | D | D |
| chr1 | TSHB:NM_001277991:exon1:c.T161C:p.V54A,TSHB:NM_000549:exon3:c.T296C:p.V99A | rs138267022 | 0,4500% | 0,3195% | 0,0900% | D | P | D |
| chr1 | TTLL10:NM_153254:exon2:c.T263C:p.I88T,TTLL10:NM_001130045:exon6:c.T482C:p.I161T | rs202096060 | 0,0080% | 0,0599% | 0,0100% | D | D | D |
| chr1 | ZNF687:NM_001304764:exon4:c.G2323A:p.G775S,ZNF687:NM_020832:exon4:c.G2323A:p.G775S,ZNF687:NM_001304763:exon5:c.G2323A:p.G775S | rs147334222 | 0,0077% | 0,0200% | 0,0037% | T | D | D |
| chr10 | BTRC:NM_001256856:exon9:c.A1162G:p.T388A,BTRC:NM_003939:exon9:c.A1132G:p.T378A,BTRC:NM_033637:exon10:c.A1240G:p.T414A | rs141008374 | 0,8000% | 0,6989% | 0,1600% | T | B | D |
| chr10 | CCDC186:NM_018017:exon2:c.G416A:p.S139N,CCDC186:NM_001321829:exon3:c.G416A:p.S139N,CCDC186:NM_153249:exon3:c.G416A:p.S139N | rs149190193 | 0,1800% | 0,0998% | 0,0300% | T | P | D |
| chr10 | H2AFY2:NM_018649:exon9:c.G1078A:p.G360S | rs1012386464 | 0,0000% | 0,0000% | 0,0008% | D | D | D |
| chr10 | HABP2:NM_001177660:exon3:c.A70T:p.N24Y,HABP2:NM_004132:exon3:c.A148T:p.N50Y | rs113225892 | 0,0200% | 0,0399% | 0,0077% | D | P | N |
| chr10 | TDRD1:NM_198795:exon4:c.T401G:p.V134G | rs746213162 | 0,0000% | 0,0000% | 0,0004% | D | P | N |
| chr11 | CCDC73:NM_001008391:exon15:c.C1252T:p.Q418X | rs199824803 | 0,0000% | 0,0799% | 0,0052% | NA | NA | A |
| chr11 | CDC42BPG:NM_017525:exon8:c.C1069T:p.R357W | rs761133866 | 0,0000% | 0,0000% | 0,0012% | D | D | N |
| chr11 | CTSW:NM_001335:exon4:c.G317A:p.R106Q | rs114953123 | 0,0800% | 0,0200% | 0,0800% | T | B | N |
| chr11 | ESRRA:NM_001282450:exon2:c.C92T:p.T31I,ESRRA:NM_001282451:exon2:c.C92T:p.T31I,ESRRA:NM_004451:exon2:c.C92T:p.T31I | rs117285599 | 0,4600% | 0,3594% | 0,3400% | T | B | N |
| chr11 | MRGPRX4:NM_054032:exon1:c.C449T:p.S150F | rs73434269 | 0,9600% | 1,0383% | 0,2100% | D | D | N |
| chr11 | PCSK7:NM_004716:exon17:c.T2051C:p.L684P | NA | 0,0000% | 0,0000% | NA | D | D | D |
| chr11 | SPI1:NM_001080547:exon5:c.C626T:p.A209V,SPI1:NM_003120:exon5:c.C623T:p.A208V | NA | 0,0000% | 0,0000% | NA | T | B | D |
| chr11 | USH1C:NM_001297764:exon11:c.G889C:p.E297Q,USH1C:NM_005709:exon12:c.G946C:p.E316Q,USH1C:NM_153676:exon12:c.G946C:p.E316Q | rs35336155 | 0,1600% | 0,5990% | 0,0900% | D | D | D |
| chr12 | ACACB:NM_001093:exon5:c.1055_1056insCCA:p.G352delinsGH | NA | 0,0000% | 0,0000% | NA | NA | NA | NA |
| chr12 | CD163L1:NM_001297650:exon13:c.G3193A:p.G1065S,CD163L1:NM_174941:exon13:c.G3163A:p.G1055S | rs36206713 | 0,6200% | 0,2396% | 0,6500% | D | D | D |
| chr12 | CLSTN3:NM_014718:exon10:c.A1500T:p.K500N | rs143003914 | 0,0400% | 0,0399% | 0,0200% | T | P | D |
| chr12 | DENND5B:NM_144973:exon3:c.T749C:p.V250A,DENND5B:NM_001308339:exon5:c.T854C:p.V285A | rs189201166 | 0,0200% | 0,0200% | 0,0026% | T | B | D |
| chr12 | GNPTAB:NM_024312:exon16:c.C3197T:p.T1066M | rs34083392 | 0,7800% | 1,0783% | 0,1800% | D | D | D |
| chr12 | GXYLT1:NM_001099650:exon6:c.G970A:p.A324T,GXYLT1:NM_173601:exon7:c.G1063A:p.A355T | rs374749531 | 0,0300% | 0,0000% | 0,0045% | D | D | D |
| chr12 | HAL:NM_001258333:exon17:c.G1021A:p.V341M,HAL:NM_001258334:exon18:c.G1645A:p.V549M,HAL:NM_002108:exon18:c.G1645A:p.V549M | rs61937878 | 0,3700% | 0,0998% | 0,2100% | D | D | D |
| chr12 | HELB:NM_033647:exon4:c.G1507T:p.A503S | rs771324513 | 0,0000% | 0,0000% | 0,0004% | D | D | D |
| chr12 | IGFBP6:NM_002178:exon4:c.G650A:p.R217Q | rs6413498 | 0,7800% | 0,7987% | 0,1800% | T | D | N |
| chr12 | MANSC1:NM_018050:exon4:c.A593G:p.Y198C | rs138574718 | 0,0500% | 0,0200% | 0,0081% | D | P | N |
| chr12 | MMAB:NM_052845:exon5:c.G403A:p.A135T | rs35648932 | 0,2500% | 0,2995% | 0,0700% | T | P | D |
| chr12 | MYBPC1:NM_001254718:exon30:c.C3497T:p.S1166F | rs73390504 | 0,0000% | 1,5974% | 0,2800% | D | P | D |
| chr12 | SLC48A1:NM_017842:exon1:c.C107T:p.P36L | rs748734372 | 0,0000% | 0,0000% | 0,0100% | T | D | D |
| chr12 | TAS2R20:NM_176889:exon1:c.A808G:p.I270V | rs138274249 | 0,3600% | 0,4193% | 0,1000% | T | B | N |
| chr12 | VEZT:NM_017599:exon6:c.A749G:p.H250R | NA | 0,0000% | 0,0000% | 0,0004% | T | P | D |
| chr13 | DIAPH3:NM_001258370:exon14:c.A1565G:p.Q522R,DIAPH3:NM_030932:exon14:c.A1565G:p.Q522R,DIAPH3:NM_001258367:exon18:c.A2216G:p.Q739R,DIAPH3:NM_001258368:exon18:c.A2144G:p.Q715R,DIAPH3:NM_001258366:exon19:c.A2321G:p.Q774R,DIAPH3:NM_001042517:exon20:c.A2354G:p.Q785R,DIAPH3:NM_001258369:exon20:c.A2354G:p.Q785R | rs199786163 | 0,0300% | 0,0399% | 0,0500% | T | P | D |
| chr13 | GPC6:NM_005708:exon3:c.A710G:p.K237R | rs201337600 | 0,0400% | 0,0599% | 0,0500% | T | B | D |
| chr14 | DICER1:NM_001195573:exon4:c.G485A:p.G162D,DICER1:NM_001271282:exon5:c.G485A:p.G162D,DICER1:NM_001291628:exon5:c.G485A:p.G162D,DICER1:NM_177438:exon5:c.G485A:p.G162D,DICER1:NM_030621:exon7:c.G485A:p.G162D | rs142815547 | 0,2100% | 0,1597% | 0,0600% | T | B | D |
| chr14 | FBXO33:NM_203301:exon1:c.9_11del:p.3_4del | rs773817973 | 0,0000% | 0,0000% | 0,0048% | NA | NA | NA |
| chr14 | GEMIN2:NM_001009182:exon5:c.G469A:p.G157R,GEMIN2:NM_001009183:exon5:c.G469A:p.G157R,GEMIN2:NM_003616:exon5:c.G469A:p.G157R | rs140431922 | 0,0600% | 0,0399% | 0,0100% | T | P | D |
| chr14 | CEP170B:NM_015005:exon7:c.C566T:p.A189V,CEP170B:NM_001112726:exon8:c.C776T:p.A259V | rs41304371 | 0,8400% | 0,8187% | 0,2600% | T | B | N |
| chr14 | KIAA0391:NM_001256678:exon2:c.C963A:p.H321Q,KIAA0391:NM_001256679:exon2:c.C678A:p.H226Q,KIAA0391:NM_014672:exon2:c.C963A:p.H321Q | rs372368846 | 0,0300% | 0,0000% | 0,0200% | T | B | D |
| chr14 | KLHL28:NM_001308112:exon3:c.G1330A:p.V444I,KLHL28:NM_017658:exon3:c.G1288A:p.V430I | rs34373760 | 0,3500% | 0,3994% | 0,0800% | D | D | D |
| chr14 | KTN1:NM_001079521:exon25:c.A2597G:p.Q866R,KTN1:NM_001079522:exon25:c.A2528G:p.Q843R,KTN1:NM_004986:exon25:c.A2597G:p.Q866R,KTN1:NM_001271014:exon26:c.A2597G:p.Q866R | rs76902262 | 0,6700% | 0,5192% | 0,1400% | D | D | D |
| chr14 | MIS18BP1:NM_018353:exon4:c.A928G:p.T310A | rs144583290 | 0,6300% | 0,6589% | 0,1500% | T | P | N |
| chr14 | MIS18BP1:NM_018353:exon2:c.G79A:p.D27N | rs144234035 | 0,6300% | 0,6589% | 0,1500% | T | B | N |
| chr14 | NFKBIA:NM_020529:exon4:c.G581C:p.G194A | rs148656104 | 0,3700% | 0,1797% | 0,0600% | D | D | D |
| chr14 | OTX2:NM_001270525:exon3:c.G444C:p.P148P,OTX2:NM_172337:exon3:c.G420C:p.P140P,OTX2:NM_001270524:exon4:c.G420C:p.P140P,OTX2:NM_001270523:exon5:c.G420C:p.P140P,OTX2:NM_021728:exon5:c.G444C:p.P148P | rs147896150 | 0,1200% | 0,1198% | 0,0400% | NA | NA | NA |
| chr14 | PLEKHG3:NM_001308147:exon17:c.G3148A:p.V1050I | rs17180132 | 0,7500% | 0,3195% | 0,6700% | D | D | D |
| chr14 | SIPA1L1:NM_001284247:exon1:c.C166A:p.P56T,SIPA1L1:NM_001284245:exon2:c.C166A:p.P56T,SIPA1L1:NM_001284246:exon2:c.C166A:p.P56T,SIPA1L1:NM_015556:exon2:c.C166A:p.P56T | rs12884638 | 0,8700% | 0,2196% | 0,7900% | D | P | D |
| chr14 | SLIRP:NM_001267863:exon1:c.C29T:p.A10V,SLIRP:NM_001267864:exon1:c.C29T:p.A10V,SLIRP:NM_031210:exon1:c.C29T:p.A10V | rs117221257 | 0,5400% | 0,4393% | 0,6100% | T | B | N |
| chr14 | TECPR2:NM_001172631:exon13:c.C2953T:p.P985S,TECPR2:NM_014844:exon13:c.C2953T:p.P985S | rs374430400 | 0,0200% | 0,0000% | 0,0012% | T | D | D |
| chr14 | TNFAIP2:NM_006291:exon7:c.C1344A:p.S448R | rs61706103 | 0,7800% | 0,8387% | 0,1800% | T | B | N |
| chr14 | WDHD1:NM_001008396:exon9:c.A457G:p.I153V,WDHD1:NM_007086:exon10:c.A826G:p.I276V | rs115947628 | 0,4500% | 0,3794% | 0,1200% | T | B | D |
| chr16 | ANKRD11:NM_001256183:exon9:c.4475_4498del:p.1492_1500del,ANKRD11:NM_013275:exon9:c.4475_4498del:p.1492_1500del,ANKRD11:NM_001256182:exon10:c.4475_4498del:p.1492_1500del | rs534329317 | 0,0000% | 0,0000% | 0,0700% | NA | NA | NA |
| chr16 | C16orf58:NM_022744:exon1:c.286_287insCTGT:p.W96fs | NA | 0,0000% | 0,0000% | NA | NA | NA | NA |
| chr16 | CDT1:NM_030928:exon2:c.C248T:p.P83L | rs139038990 | 0,1300% | 0,1398% | 0,6900% | T | D | N |
| chr16 | HIRIP3:NM_001197323:exon4:c.G325C:p.G109R,HIRIP3:NM_003609:exon5:c.G1263C:p.E421D | rs138542330 | 0,0800% | 0,0399% | 0,0100% | D | P | D |
| chr16 | PDXDC1:NM_001285444:exon22:c.G2081A:p.S694N,PDXDC1:NM_001285445:exon22:c.G2078A:p.S693N,PDXDC1:NM_001285448:exon22:c.G1889A:p.S630N,PDXDC1:NM_001285447:exon23:c.G2117A:p.S706N,PDXDC1:NM_001324019:exon23:c.G2159A:p.S720N,PDXDC1:NM_015027:exon23:c.G2162A:p.S721N | rs148061029 | 0,3100% | 0,2596% | 0,2800% | D | D | D |
| chr16 | ROGDI:NM_024589:exon8:c.C532T:p.R178W | rs566665742 | 0,0000% | 0,0200% | 0,0029% | D | D | D |
| chr16 | SPIRE2:NM_032451:exon5:c.A889G:p.M297V | rs139065194 | 0,0077% | 0,0000% | 0,0200% | D | B | D |
| chr16 | SRCAP:NM_006662:exon34:c.A8350C:p.T2784P | rs73538429 | 0,4500% | 0,3994% | 0,0800% | T | P | N |
| chr16 | ZNF629:NM_001080417:exon3:c.G718C:p.E240Q,ZNF629:NM_001345970:exon3:c.G676C:p.E226Q | NA | 0,0000% | 0,0000% | NA | D | D | D |
| chr17 | ASB16:NM_080863:exon5:c.G1283A:p.R428Q | rs140865656 | 0,1500% | 0,1997% | 0,2600% | T | P | N |
| chr17 | FAM83G:NM_001039999:exon5:c.G1374T:p.Q458H | rs201344489 | 0,7100% | 0,1597% | 0,5600% | D | D | N |
| chr17 | IFI35:NM_001330230:exon2:c.A52G:p.R18G,IFI35:NM_005533:exon2:c.A52G:p.R18G | rs148786311 | 0,1300% | 0,1597% | 0,2500% | D | P | N |
| chr17 | IGFBP4:NM_001552:exon1:c.C121A:p.P41T | rs758124086 | 0,0000% | 0,0000% | 0,0064% | D | D | D |
| chr17 | MYH8:NM_002472:exon30:c.G4042A:p.E1348K | rs140562514 | 0,1400% | 0,0399% | 0,0700% | D | D | D |
| chr17 | PLD2:NM_001243108:exon20:c.G2057C:p.G686A,PLD2:NM_002663:exon20:c.G2057C:p.G686A | rs147534454 | 0,1200% | 0,0599% | 0,0800% | T | D | D |
| chr17 | RUNDC3A:NM_001144825:exon1:c.C23T:p.T8I,RUNDC3A:NM_001144826:exon1:c.C23T:p.T8I,RUNDC3A:NM_006695:exon1:c.C23T:p.T8I | NA | 0,0000% | 0,0000% | NA | D | B | N |
| chr17 | TMEM102:NM_001320444:exon2:c.G227T:p.R76L,TMEM102:NM_178518:exon3:c.G227T:p.R76L | rs147336927 | 0,0400% | 0,0200% | 0,0700% | D | P | D |
| chr17 | TNS4:NM_032865:exon4:c.G1192T:p.V398F | rs140876567 | 0,0000% | 0,8187% | 0,3400% | D | P | N |
| chr18 | SERPINB10:NM_005024:exon3:c.G205T:p.D69Y | rs143009235 | 0,5300% | 0,3594% | 0,5500% | T | B | N |
| chr19 | ABCA7:NM_019112:exon19:c.G2639A:p.R880Q | rs143718918 | 0,0600% | 0,0200% | 0,1100% | D | D | D |
| chr19 | CACNA1A:NM_001127221:exon19:c.C2740T:p.P914S,CACNA1A:NM_001127222:exon19:c.C2737T:p.P913S | rs16020 | 0,5800% | 0,6190% | 0,1400% | T | B | N |
| chr19 | COL5A3:NM_015719:exon49:c.T3584C:p.V1195A | rs62638750 | 0,8000% | 0,2796% | 0,8600% | T | B | N |
| chr19 | HSH2D:NM_001291274:exon9:c.A704G:p.K235R,HSH2D:NM_032855:exon9:c.A875G:p.K292R | rs199804128 | 0,4600% | 0,1198% | 0,3700% | NA | B | N |
| chr19 | LRRC25:NM_145256:exon1:c.A22G:p.T8A | rs139562741 | 0,1400% | 0,0599% | 0,2000% | T | B | N |
| chr19 | RFXANK:NM_001278728:exon2:c.C95T:p.A32V,RFXANK:NM_001278727:exon3:c.C95T:p.A32V,RFXANK:NM_003721:exon3:c.C95T:p.A32V,RFXANK:NM_134440:exon3:c.C95T:p.A32V | rs114064359 | 0,1500% | 0,1597% | 0,0300% | T | B | N |
| chr19 | SUGP2:NM_001017392:exon3:c.G386C:p.G129A,SUGP2:NM_001321697:exon3:c.G386C:p.G129A,SUGP2:NM_001321698:exon3:c.G428C:p.G143A,SUGP2:NM_001321699:exon3:c.G428C:p.G143A,SUGP2:NM_014884:exon3:c.G386C:p.G129A | rs74855957 | 0,4800% | 0,4792% | 0,1200% | D | D | N |
| chr19 | ZNF599:NM_001007248:exon4:c.G1258T:p.E420X | rs112895297 | 0,3500% | 0,0599% | 0,4800% | NA | NA | D |
| chr19 | ZNF98:NM_001098626:exon4:c.A1195G:p.I399V | NA | 0,0000% | 0,0000% | NA | D | P | N |
| chr2 | ALLC:NM_018436:exon9:c.G673T:p.G225C | rs200086179 | 0,0800% | 0,0599% | 0,1100% | D | D | D |
| chr2 | CAPN10:NM_023083:exon6:c.T901C:p.F301L,CAPN10:NM_023085:exon6:c.T901C:p.F301L | NA | 0,0000% | 0,0000% | NA | D | D | D |
| chr2 | CCDC148:NM_001301685:exon3:c.T217C:p.W73R,CCDC148:NM_138803:exon3:c.T217C:p.W73R | NA | 0,0000% | 0,0000% | NA | D | D | D |
| chr2 | CHRND:NM_001311195:exon6:c.C280G:p.Q94E,CHRND:NM_001256657:exon7:c.C817G:p.Q273E,CHRND:NM_000751:exon8:c.C862G:p.Q288E,CHRND:NM_001311196:exon8:c.C559G:p.Q187E | rs41265127 | 0,3800% | 0,0399% | 0,2700% | D | D | D |
| chr2 | DNAH6:NM_001370:exon72:c.G11669A:p.R3890H | rs200449901 | 0,0000% | 0,0399% | 0,0300% | D | D | D |
| chr2 | GALNT5:NM_014568:exon5:c.C1922T:p.P641L,GALNT5:NM_001329868:exon6:c.C512T:p.P171L | NA | 0,0000% | 0,0000% | NA | D | D | D |
| chr2 | LRP1B:NM_018557:exon86:c.C13229A:p.T4410K | rs772444623 | 0,0000% | 0,0000% | 0,0004% | T | D | D |
| chr2 | MALL:NM_005434:exon2:c.G139A:p.A47T | rs141817647 | 0,0400% | 0,0000% | 0,1900% | T | P | D |
| chr2 | PASK:NM_001252119:exon3:c.C287T:p.T96M,PASK:NM_001252122:exon3:c.C287T:p.T96M,PASK:NM_001252124:exon3:c.C287T:p.T96M,PASK:NM_015148:exon3:c.C287T:p.T96M,PASK:NM_001252120:exon4:c.C287T:p.T96M | rs778811482 | 0,0000% | 0,0000% | 0,0008% | D | D | N |
| chr2 | RGPD3:NM_001144013:exon3:c.C236T:p.A79V | rs199757971 | 0,0000% | 0,0000% | 0,4800% | D | D | D |
| chr2 | RGPD4:NM_182588:exon21:c.C5004G:p.H1668Q | rs202136029 | 0,0000% | 0,1198% | 0,2600% | T | D | N |
| chr2 | CAVIN2:NM_004657:exon2:c.C626A:p.P209H | rs34841326 | 0,3600% | 0,1597% | 0,4200% | D | B | N |
| chr2 | SNED1:NM_001080437:exon13:c.C1768T:p.R590W | NA | 0,0000% | 0,0000% | 0,0013% | T | D | D |
| chr2 | STK36:NM_001243313:exon26:c.G3563A:p.R1188Q,STK36:NM_015690:exon26:c.G3626A:p.R1209Q | rs201010018 | 0,0077% | 0,0000% | 0,0200% | D | D | D |
| chr2 | ZRANB3:NM_001286568:exon9:c.C1013T:p.S338L,ZRANB3:NM_032143:exon9:c.C1013T:p.S338L | rs61733463 | 0,1800% | 0,3195% | 0,1000% | T | B | N |
| chr2 | C20orf196:NM_152504:exon3:c.C427T:p.R143C,C20orf196:NM_001303477:exon4:c.C427T:p.R143C,C20orf196:NM_001303478:exon4:c.C331T:p.R111C | rs74824950 | 0,5900% | 0,3794% | 0,1700% | D | D | D |
| chr20 | SLX4IP:NM_001009608:exon5:c.C314T:p.A105V | rs140984471 | 0,5800% | 0,1398% | 0,5700% | T | B | N |
| chr20 | NCOA3:NM_001174087:exon15:c.C2810G:p.S937C,NCOA3:NM_001174088:exon15:c.C2795G:p.S932C,NCOA3:NM_006534:exon15:c.C2810G:p.S937C,NCOA3:NM_181659:exon15:c.C2810G:p.S937C | rs142951578 | 0,0500% | 0,0998% | 0,0400% | D | P | D |
| chr21 | KRTAP20-2:NM_181616:exon1:c.G155A:p.R52H | rs760841338 | 0,0000% | 0,0000% | 0,0041% | NA | B | N |
| chr21 | MX1:NM_001282920:exon4:c.T512C:p.V171A,MX1:NM_001178046:exon6:c.T512C:p.V171A,MX1:NM_002462:exon8:c.T512C:p.V171A,MX1:NM_001144925:exon10:c.T512C:p.V171A | rs951248333 | 0,0000% | 0,0000% | NA | D | P | D |
| chr21 | TFF2:NM_005423:exon1:c.C7T:p.R3W | rs7277409 | 0,8500% | 0,6190% | 0,1700% | T | B | N |
| chr21 | TRPM2:NM_001320352:exon4:c.G156T:p.M52I,TRPM2:NM_001320351:exon28:c.G4011T:p.M1337I,TRPM2:NM_003307:exon29:c.G4113T:p.M1371I,TRPM2:NM_001320350:exon30:c.G4263T:p.M1421I | NA | 0,0000% | 0,0000% | NA | T | B | D |
| chr21 | UBASH3A:NM_001001895:exon4:c.G502A:p.V168I,UBASH3A:NM_001243467:exon4:c.G502A:p.V168I,UBASH3A:NM_018961:exon4:c.G502A:p.V168I | rs139138633 | 0,1800% | 0,1797% | 0,0400% | T | B | N |
| chr21 | ZBTB21:NM_001320729:exon2:c.1416_1418del:p.472_473del,ZBTB21:NM_020727:exon2:c.1416_1418del:p.472_473del,ZBTB21:NM_001098402:exon3:c.1416_1418del:p.472_473del,ZBTB21:NM_001098403:exon3:c.1416_1418del:p.472_473del,ZBTB21:NM_001320731:exon4:c.1416_1418del:p.472_473del | rs200509586 | 0,0200% | 0,7388% | 0,6700% | NA | NA | NA |
| chr22 | BRD1:NM_001349941:exon7:c.C2830T:p.R944C,BRD1:NM_001304808:exon8:c.C2845T:p.R949C,BRD1:NM_001304809:exon8:c.C2452T:p.R818C,BRD1:NM_001349940:exon8:c.C2599T:p.R867C,BRD1:NM_001349942:exon8:c.C1303T:p.R435C | rs139630692 | 0,0700% | 0,0399% | 0,0500% | D | D | D |
| chr22 | HMGXB4:NM_001003681:exon5:c.G558T:p.E186D | rs41282603 | 0,0000% | 0,0200% | 0,0200% | D | D | D |
| chr22 | IL2RB:NM_000878:exon4:c.G235A:p.V79M,IL2RB:NM_001346222:exon4:c.G235A:p.V79M,IL2RB:NM_001346223:exon4:c.G235A:p.V79M | rs149508414 | 0,2200% | 0,1997% | 0,3600% | T | B | N |
| chr22 | SBF1:NM_002972:exon37:c.C5047G:p.L1683V | rs776198448 | 0,0000% | 0,0000% | 0,0017% | T | B | D |
| chr3 | EEFSEC:NM_021937:exon5:c.G1040A:p.R347Q | rs149382419 | 0,0800% | 0,0000% | 0,0500% | T | B | D |
| chr3 | ITPR1:NM_002222:exon39:c.G5146A:p.G1716S,ITPR1:NM_001099952:exon40:c.G5191A:p.G1731S,ITPR1:NM_001168272:exon42:c.G5290A:p.G1764S | rs369108656 | 0,0082% | 0,0000% | 0,0100% | T | B | D |
| chr4 | RGS12:NM_002926:exon2:c.G302A:p.R101H,RGS12:NM_198229:exon2:c.G302A:p.R101H | rs75169970 | 0,6700% | 0,9784% | 0,1700% | T | B | N |
| chr4 | SOWAHB:NM_001029870:exon1:c.G1678A:p.E560K | rs141644004 | 0,1800% | 0,2396% | 0,0500% | D | B | N |
| chr4 | UGT2B28:NM_001207004:exon1:c.G4A:p.A2T,UGT2B28:NM_053039:exon1:c.G4A:p.A2T | rs142644303 | 0,7200% | 0,3794% | 0,1300% | T | B | N |
| chr4 | USP38:NM_001290325:exon9:c.C1795T:p.R599C,USP38:NM_032557:exon9:c.C1795T:p.R599C,USP38:NM_001290326:exon10:c.C430T:p.R144C | rs75962202 | 0,8800% | 0,8187% | 0,1700% | T | B | N |
| chr4 | ZNF141:NM_001348277:exon2:c.C533G:p.P178R,ZNF141:NM_003441:exon4:c.C761G:p.P254R | rs782497343 | 0,0000% | 0,0000% | 0,0016% | D | D | D |
| chr4 | ZNF141:NM_001348277:exon2:c.T787G:p.Y263D,ZNF141:NM_003441:exon4:c.T1015G:p.Y339D | rs372553337 | 0,0077% | 0,0000% | 0,0016% | D | D | N |
| chr5 | DDX4:NM_001166534:exon11:c.G1108A:p.V370I,DDX4:NM_001142549:exon17:c.G1453A:p.V485I,DDX4:NM_001166533:exon17:c.G1495A:p.V499I,DDX4:NM_024415:exon18:c.G1555A:p.V519I | rs146056716 | 0,0077% | 0,0000% | 0,0100% | T | B | N |
| chr5 | NA | NA | 0,0000% | 0,0000% | NA | NA | NA | NA |
| chr5 | FYB:NM_001243093:exon2:c.A1025G:p.K342R,FYB:NM_001465:exon2:c.A995G:p.K332R,FYB:NM_199335:exon2:c.A995G:p.K332R,FYB:NM_001349333:exon3:c.A995G:p.K332R | rs3749741 | 0,0083% | 0,8786% | 0,4800% | T | B | N |
| chr5 | PCDHB15:NM_018935:exon1:c.T1561C:p.Y521H | rs148237600 | 0,3200% | 0,0599% | 0,1600% | D | D | D |
| chr5 | ARHGEF28:NM_001244364:exon8:c.A1013G:p.N338S,ARHGEF28:NM_001080479:exon16:c.A1952G:p.N651S,ARHGEF28:NM_001177693:exon16:c.A1952G:p.N651S | rs187509753 | 0,2400% | 0,0599% | 0,4900% | T | B | N |
| chr5 | TRIO:NM_007118:exon55:c.A8543C:p.Q2848P | NA | 0,0000% | 0,0000% | NA | D | D | D |
| chr6 | CRYBG1:NM_001624:exon10:c.T3909A:p.N1303K | rs144800958 | 0,0200% | 0,0200% | 0,0200% | D | B | D |
| chr6 | AK9:NM_001145128:exon12:c.C1231G:p.L411V,AK9:NM_001329602:exon12:c.C1231G:p.L411V,AK9:NM_145025:exon12:c.C1231G:p.L411V,AK9:NM_001329603:exon13:c.C1447G:p.L483V | NA | 0,0000% | 0,0000% | NA | D | P | D |
| chr6 | CASP8AP2:NM_001137667:exon8:c.A2750T:p.D917V,CASP8AP2:NM_001137668:exon8:c.A2750T:p.D917V,CASP8AP2:NM_012115:exon8:c.A2750T:p.D917V | rs199818755 | 0,0300% | 0,0000% | 0,0055% | NA | P | NA |
| chr6 | COL21A1:NM_001318752:exon4:c.G677A:p.R226H,COL21A1:NM_030820:exon4:c.G677A:p.R226H,COL21A1:NM_001318751:exon5:c.G677A:p.R226H | rs572301223 | 0,0000% | 0,0599% | 0,0400% | D | D | D |
| chr6 | FAM184A:NM_001100411:exon7:c.G1352A:p.G451E,FAM184A:NM_001288576:exon7:c.G1352A:p.G451E,FAM184A:NM_024581:exon7:c.G1712A:p.G571E | NA | 0,0000% | 0,0000% | NA | D | D | D |
| chr6 | GJA10:NM_032602:exon1:c.G1390A:p.E464K | rs140533217 | 0,0800% | 0,0399% | 0,1200% | D | B | N |
| chr6 | RSPH9:NM_001193341:exon1:c.G146T:p.R49L,RSPH9:NM_152732:exon1:c.G146T:p.R49L | rs1055513498 | 0,0000% | 0,0000% | NA | D | P | D |
| chr7 | FAM71F1:NM_001282788:exon3:c.C538T:p.P180S,FAM71F1:NM_032599:exon3:c.C538T:p.P180S,FAM71F1:NM_001282789:exon4:c.C241T:p.P81S | rs371977508 | 0,0077% | 0,0000% | 0,0004% | D | P | N |
| chr7 | GPNMB:NM_001005340:exon11:c.G1699T:p.E567X,GPNMB:NM_002510:exon11:c.G1663T:p.E555X | rs11537976 | 0,3600% | 0,4193% | 1,0200% | NA | NA | D |
| chr7 | HOXA13:NM_000522:exon1:c.G124A:p.A42T | NA | 0,0000% | 0,0000% | NA | D | P | N |
| chr7 | KIAA1549:NM_001164665:exon6:c.G3317A:p.R1106Q,KIAA1549:NM_020910:exon6:c.G3317A:p.R1106Q | rs200423257 | 0,1100% | 0,0200% | 0,0700% | T | D | N |
| chr7 | MGAM:NM_004668:exon9:c.C1018T:p.R340C | rs199961575 | 0,0700% | 0,0399% | 0,0600% | D | D | D |
| chr7 | MICALL2:NM_182924:exon6:c.G1150A:p.V384M | rs202171352 | 0,1200% | 0,0000% | 0,1100% | T | D | N |
| chr7 | NOM1:NM_138400:exon7:c.A1946T:p.Q649L | rs114916388 | 0,0300% | 0,0399% | 0,0100% | D | B | D |
| chr7 | RABGEF1:NM_001287061:exon3:c.T248A:p.F83Y,RABGEF1:NM_014504:exon3:c.T206A:p.F69Y,RABGEF1:NM_001287062:exon4:c.T206A:p.F69Y | NA | 0,0000% | 0,0000% | NA | T | B | D |
| chr8 | CA2:NM_000067:exon2:c.C102A:p.D34E | rs201063453 | 0,0000% | 0,0000% | 0,0004% | T | B | N |
| chr8 | COL14A1:NM_021110:exon21:c.C2530T:p.R844W | rs115276090 | 0,2800% | 0,1398% | 0,1900% | D | D | D |
| chr8 | FAM160B2:NM_022749:exon15:c.G1892A:p.R631Q | rs113817175 | 0,5300% | 0,3395% | 0,5900% | D | D | D |
| chr8 | MTBP:NM_022045:exon19:c.C2248T:p.L750F | rs61753798 | 0,8200% | 0,5391% | 0,7700% | T | B | D |
| chr8 | MTERF3:NM_001286643:exon3:c.C353A:p.P118Q,MTERF3:NM_015942:exon3:c.C353A:p.P118Q | rs75477527 | 0,8800% | 0,4193% | 0,8400% | T | D | D |
| chr8 | SLC45A4:NM_001080431:exon4:c.G946A:p.A316T,SLC45A4:NM_001286646:exon4:c.G1099A:p.A367T,SLC45A4:NM_001286648:exon4:c.G946A:p.A316T | rs61995887 | 0,2800% | 0,2396% | 0,0500% | T | B | N |
| chr8 | SLC45A4:NM_001080431:exon2:c.G247A:p.V83I,SLC45A4:NM_001286646:exon2:c.G400A:p.V134I,SLC45A4:NM_001286648:exon2:c.G247A:p.V83I | rs150530787 | 0,0400% | 0,0200% | 0,0081% | T | P | D |
| chr8 | SPATC1:NM_001134374:exon2:c.G464A:p.S155N,SPATC1:NM_198572:exon2:c.G464A:p.S155N | rs138478557 | 0,3900% | 0,1398% | 0,3200% | T | D | N |
| chr9 | FRMD3:NM_001244962:exon2:c.C529T:p.R177W,FRMD3:NM_001244961:exon9:c.C976T:p.R326W,FRMD3:NM_001244959:exon14:c.C1558T:p.R520W,FRMD3:NM_001244960:exon14:c.C1426T:p.R476W,FRMD3:NM_174938:exon14:c.C1558T:p.R520W | rs190298518 | 0,1600% | 0,2196% | 0,1700% | T | B | D |
| chr9 | ODF2:NM_001351581:exon20:c.G2600A:p.R867H,ODF2:NM_001351587:exon20:c.G2282A:p.R761H,ODF2:NM_001351588:exon20:c.G2225A:p.R742H,ODF2:NM_001242352:exon21:c.G2453A:p.R818H,ODF2:NM_001242353:exon21:c.G2468A:p.R823H,ODF2:NM_001351577:exon21:c.G2873A:p.R958H,ODF2:NM_001351578:exon21:c.G2717A:p.R906H,ODF2:NM_001351579:exon21:c.G2609A:p.R870H,ODF2:NM_001351583:exon21:c.G2552A:p.R851H,ODF2:NM_001351584:exon21:c.G2453A:p.R818H,ODF2:NM_001351586:exon21:c.G2396A:p.R799H,ODF2:NM_153433:exon21:c.G2468A:p.R823H,ODF2:NM_153435:exon21:c.G2660A:p.R887H,ODF2:NM_001351580:exon22:c.G2609A:p.R870H,ODF2:NM_001351582:exon22:c.G2585A:p.R862H,ODF2:NM_001351585:exon22:c.G2453A:p.R818H,ODF2:NM_002540:exon23:c.G2396A:p.R799H | rs142129915 | 0,6000% | 0,2596% | 0,6000% | T | P | D |
| chr9 | SPATA6L:NM_001039395:exon5:c.C389G:p.S130C | rs188220301 | 0,1900% | 0,1398% | 0,0400% | D | B | D |
| chr9 | SPATA6L:NM_001039395:exon3:c.C97T:p.L33F | rs370244730 | 0,0000% | 0,0000% | 0,0053% | T | P | D |
| chr9 | SYK:NM_001135052:exon5:c.A784C:p.I262L,SYK:NM_001174167:exon5:c.A784C:p.I262L,SYK:NM_001174168:exon5:c.A784C:p.I262L,SYK:NM_003177:exon5:c.A784C:p.I262L | rs112619650 | 0,5900% | 0,4593% | 0,1800% | T | B | N |
| chr9 | TGFBR1:NM_001130916:exon3:c.T557C:p.I186T,TGFBR1:NM_001306210:exon4:c.T800C:p.I267T,TGFBR1:NM_004612:exon4:c.T788C:p.I263T | NA | 0,0000% | 0,0000% | NA | D | D | D |
