## Supplementary figures and images for "*NCOA3* identified as a new candidate to explain autosomal dominant progressive hearing loss"

### SUPPLEMENTAL FIGURE 1

Supplementary Figure 1

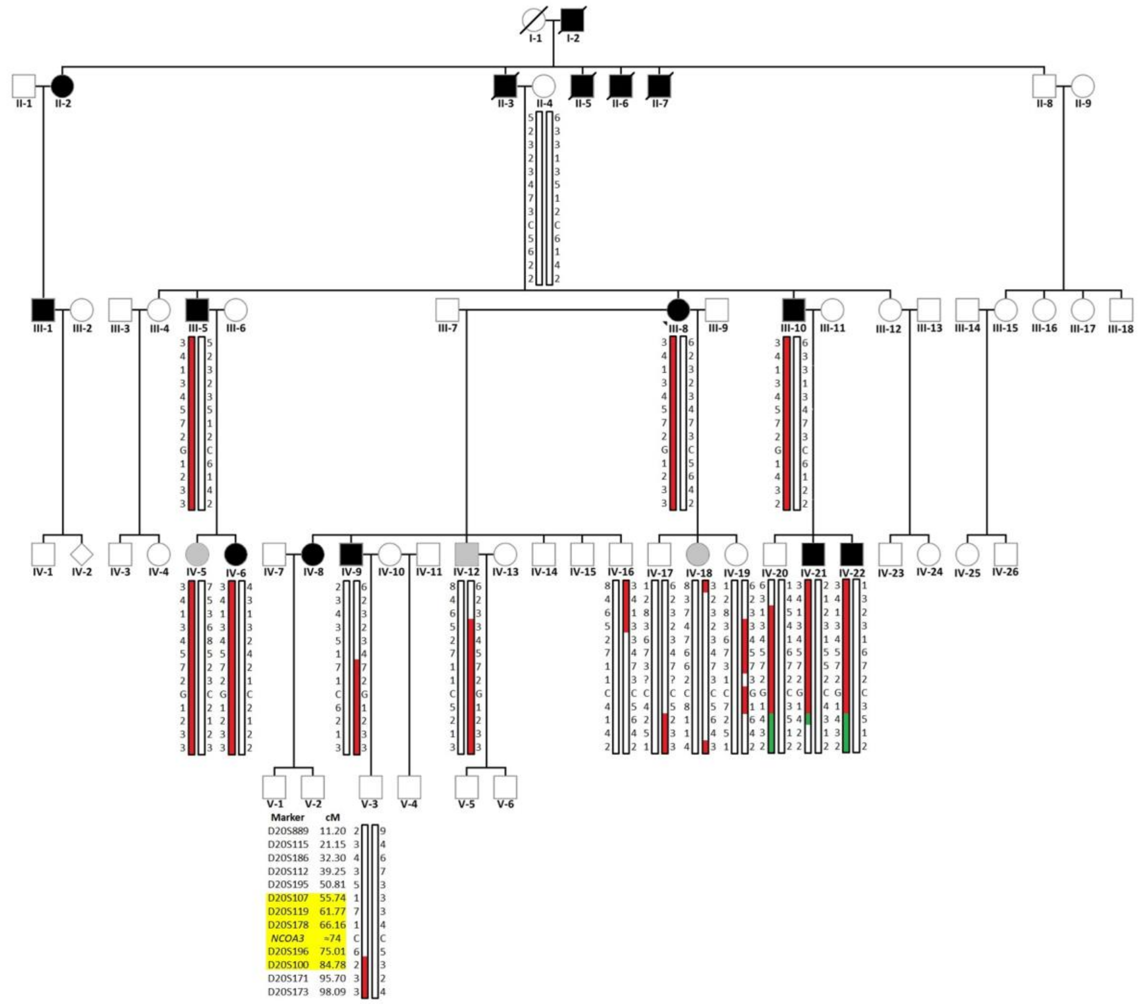

### SUPPLEMENTAL FIGURE 2

Supplementary Figure 2

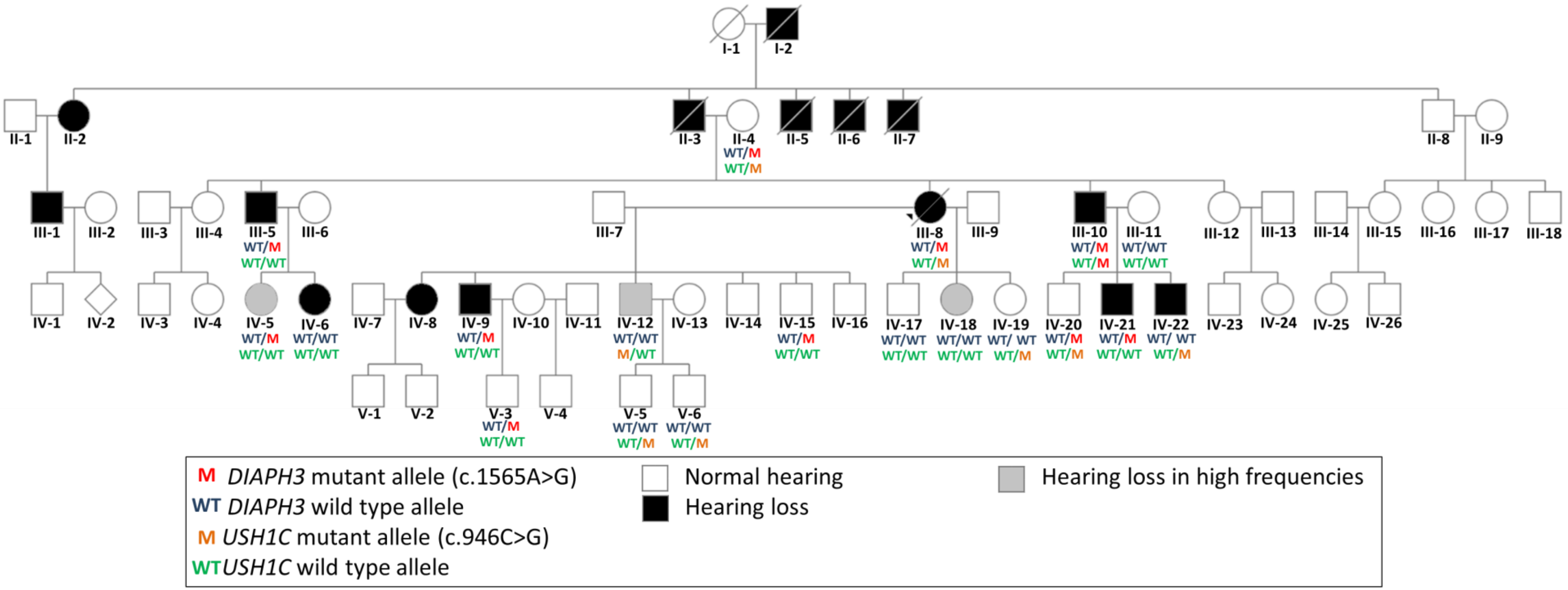

### SUPPLEMENTAL FIGURE 3

Supplementary Figure 3

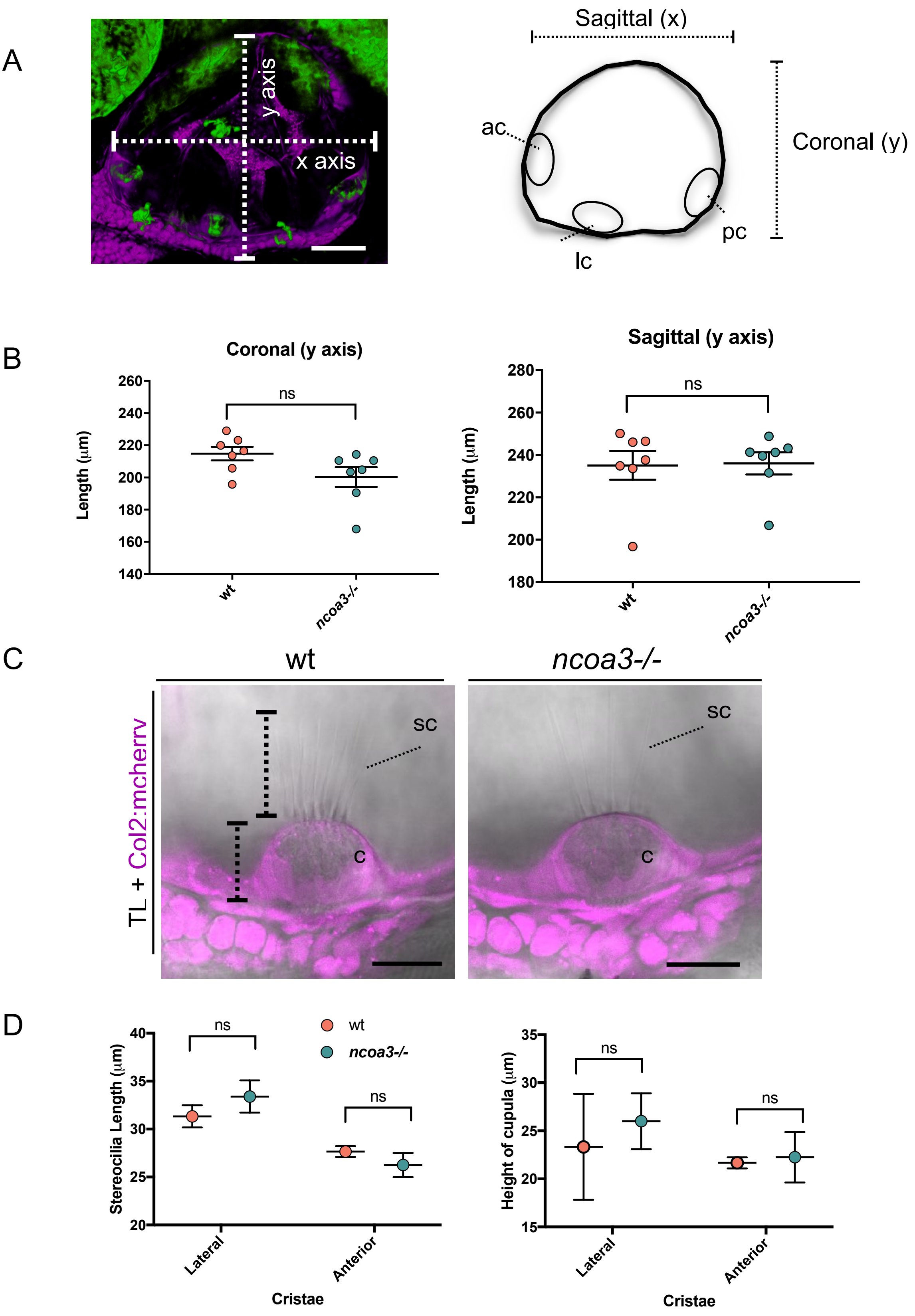

### SUPPLEMENTAL FIGURE 4

Supplementary Figure 4

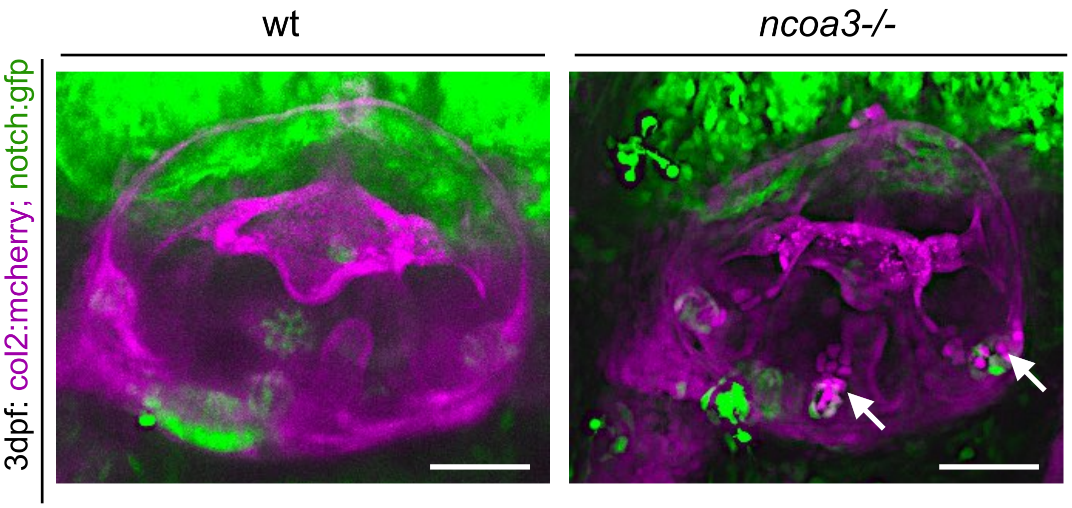
